# Nodavirus RNA Replication Crown Architecture Reveals Proto-Crown Precursor and Viral Protein A Conformational Switching

**DOI:** 10.1101/2022.12.16.520638

**Authors:** Hong Zhan, Nuruddin Unchwaniwala, Andrea Rebolledo-Viveros, Janice Pennington, Mark Horswill, Roma Broadberry, Jonathan Myers, Johan A. den Boon, Timothy Grant, Paul Ahlquist

## Abstract

Positive-strand RNA viruses replicate their genomes in virus-induced membrane vesicles, and the resulting RNA replication complexes are a major target for virus control. Nodavirus studies first revealed viral RNA replication proteins forming a 12-fold symmetric “crown” at the vesicle opening to the cytosol, an arrangement recently confirmed to extend to distantly related alphaviruses. Using cryo-electron microscopy (cryo-EM), we show that mature nodavirus crowns comprise two stacked 12-mer rings of multi-domain viral RNA replication protein A. Each ring contains an ^~^19 nm circle of C-proximal polymerase domains, differentiated by strikingly diverged positions of N-proximal RNA capping/membrane binding domains. The lower ring is a “proto-crown” precursor that assembles prior to RNA template recruitment, RNA synthesis and replication vesicle formation. In this proto-crown, the N-proximal segments interact to form a toroidal central floor, whose 3.1 Å resolution structure reveals many mechanistic details of the RNA capping/membrane binding domains. In the upper ring, cryo-EM fitting indicates that the N-proximal domains extend radially outside the polymerases, forming separated, membrane-binding “legs.” The polymerase and N-proximal domains are connected by a long linker accommodating the conformational switch between the two rings and possibly also polymerase movements associated with RNA synthesis and non-symmetric electron density in the lower center of mature crowns. The results reveal remarkable viral protein multifunctionality, conformational flexibility and evolutionary plasticity and new insights into (+)RNA virus replication and control.

**Significance:** Positive-strand RNA viruses - including coronaviruses, alphaviruses, flaviviruses and many other medically and economically important pathogens - replicate their RNA genomes by virus-encoded machinery that has been poorly characterized. Using an advanced nodavirus model, we identify a major precursor in RNA replication complex assembly and show it to be a 12-mer ring of viral RNA replication protein A, whose single particle cryo-EM structure reveals functional features of its membrane interaction, assembly, polymerase and RNA capping domains. We further show that fully functional RNA replication complexes acquire a second 12-mer ring of protein A in alternate conformation atop the first, and a central density likely to represent another polymerase conformation. These findings provide strong foundations for understanding, controlling and beneficially using such viruses.

## Introduction

Positive-strand RNA ((+)RNA) viruses include many established and emerging pathogens such as coronaviruses, flaviviruses, the alphavirus- and picornavirus-like superfamilies and numerous others. These viruses reproduce their genomes through RNA intermediates (Fig. 1A) in membrane-bounded, organelle-like RNA replication complexes (RC’s) that organize RNA replication factors and templates, protect dsRNA replication intermediates from innate immune recognition and responses, and release new (+)RNA genomes for translation, further replication and encapsidation. The two known RC classes are ^~^50-150 nm spherular invaginations or spherules, and ^~^200-300 nm double membrane vesicles or DMVs (1).

**Figure 1.**
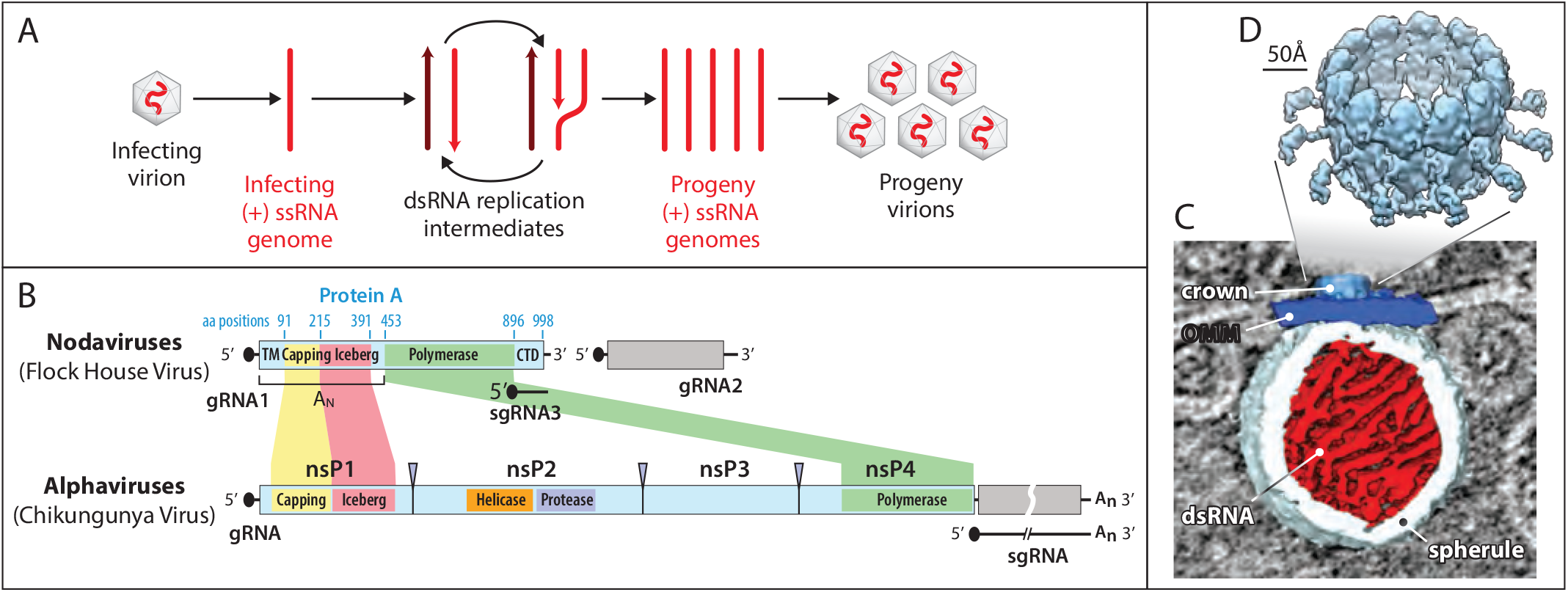
Nodavirus genome replication overview. **(A)** General mechanism of RNA genome replication by (+) strand RNA viruses. Incoming (+) RNA is used as a template for (-) strand synthesis, yielding a dsRNA replicative intermediate that templates synthesis of many copies of (+) strand genomic RNA for packaging in progeny virions. **(B)** The bipartite FHV RNA genome encodes RNA replication protein A on RNA1 (3.1 kb) and the capsid protein precursor on RNA2 (1.4 kb). Subgenomic RNA3, transcribed from RNA1, encodes RNAi suppressor B2 and an overlapping ORF B1, comprising the C-terminal 102 aa of protein A. Aligning FHV protein A and the alphavirus RNA replication polyprotein highlights conserved RNA capping and polymerase domains and FHV’s lack of helicase and protease domains. The N-terminal half of protein A, denoted segment A_N_, and alphavirus nsP1 show particular parallels. **(C)** Cryo-ET imaging of a nodavirus RC spherule in infected Drosophila S2 cells with the mitochondrial outer membrane in dark blue, invaginated spherule membrane in white, interior spherule fibrils corresponding to viral dsRNA in red, and the protein A-containing crown complex in light blue. **(D)** Reconstruction of the 12-fold symmetric nodavirus RC crown by subtomogram averaging (10).

One well-studied model for spherule RC structure and function is flock house virus (FHV), best-studied member of the nodaviruses, which infect vertebrates or invertebrates (2). FHV virions infect *Drosophila* and other insect cells,and contain a 4.5 kb genome divided between RNA1 and RNA2 (Fig. 1B). Despite its small size, the transfected FHV genome directs RNA replication and virion production in cells from invertebrates, vertebrates, yeast and plants (3, 4). This small genome size, wide replicative competence and ribosomal RNA-level viral RNA yields make FHV a potentially valuable platform for engineered RNA replication.

FHV genomic RNA1 encodes protein A, the sole viral protein required for RNA replication, which contains membrane interaction domains, a methyltransferase / guanylyltransferase (MTase/GTase) RNA capping domain, an “Iceberg” domain, and a C-proximal RNA-dependent RNA polymerase (Pol) domain (Fig. 1B). These domains are conserved throughout the distantly related alphavirus-like superfamily of viruses (5), which additionally preserve an NTPase/RNA helicase domain lacking in nodaviruses (Fig. 1B). In particular, the N-terminal 452 amino acids (aa) of protein A, hereafter denoted segment A_N_, include the MTase-GTase and Iceberg domains and parallel the RNA capping and membrane binding functions of the nsP1 proteins of classical alphaviruses like chikungunya virus (CHIKV) (Fig. 1B) (5, 6).

FHV protein A binds outer mitochondrial membranes (OMMs) and induces ^~^30-70 nm spherule invaginations (7, 8). Cryo-electron tomography (cryo-ET) of these nodavirus RCs visualized the dsRNA genome replication intermediates inside spherules and revealed that viral RNA replication protein A was localized in a striking 12-fold symmetric ring, dubbed the crown, on the cytosolic side of the spherule neck (Fig. 1C-D) (9). Further improvements resolved the major crown features (Fig. 1D) including an ^~^19 nm diameter central turret composed of 12 pairs of stacked apical and basal lobes, an inner toroidal floor, and legs projecting outward from the basal lobes, and showed by genetic tagging that each apical domain represents a protein A Pol domain (10). Protein A is the sole FHV protein required to form full crowns indistinguishable from those in FHV-infected cells (11).

Recent results imply that similar crown-like complexes of viral RNA replication proteins play crucial roles in the RCs of many, if not most, (+)RNA viruses ((1, 10, 12) and references therein). CHIKV nsP1, which as noted above is related to protein A’s A_N_ segment (Fig. 1B) (5), assembles into a 12-fold symmetric ring similar to the base of the nodavirus crown (13, 14). At even greater evolutionary distance, the double membrane vesicle (DMV) RCs of coronaviruses bear ring-shaped, viral nsp3-containing crown complexes also providing entry and exit portals to the viral RNA-rich DMV interior (15, 16).

Nodavirus RCs assemble through distinct, sequential states. In the absence of other viral components, protein A targets itself to OMMs (17, 18). Protein A specifically recognizes viral RNA templates and recruits them to the OMM (18, 19), then synthesizes (-)RNA and captures the resulting dsRNA product in an RC vesicle newly invaginated on the OMM (20). When trapped on OMMs in its pre-RNA synthesis state by polymerase mutation or absence of a replicable FHV RNA template, protein A fails to induce RNA replication vesicles and causes extensive, close zippering of adjacent mitochondria (20). The spaces between such zippered mitochondria contain regular OMM-linked structures with ^~^20 nm periodicity (20). Thus, prior to RNA synthesis, protein A induces OMM-linked complexes with dimensions similar to the mature crowns of active RCs, but with dramatically different interaction properties.

Here we show that, prior to RNA replication, protein A initially assembles on OMMs in a 12-mer ring corresponding to the basal lobe ring and floor of the mature, RNA replication-active crown. The single-particle cryo-EM structure of this proto-crown precursor shows that the A_N_ segments form the floor. The MTase-GTase and a central *α*-helical bundle share structural similarities to CHIKV nsP1 while their orientations and membrane interaction domains differ from nsP1. The Pol domain resides atop the outer edge of the A_N_ floor, in the position corresponding to the full crown basal lobes. The Pol structure fits similarly well into the full crown apical lobes, confirming prior results, and the A_N_ structure also fits well into the full crown legs. The results thus indicate that the mature crown consists of two sequentially added, stacked 12-mer rings of protein A in distinct conformations. A long, flexible linker connecting A_N_ and Pol, a Pol-like density centered in the mature crown floor, and implications for crown assembly, function and evolution are also discussed.

## Results

### Protein A alone forms “proto-crowns”

As noted in the Introduction, when nodavirus RNA replication is blocked by omitting the RNA template or mutating the template or Pol active site, protein A assembles on OMMs in ordered arrangements with interactive properties distinct from the crowns of full FHV infection (20). To determine the structure of these “pre-replication” protein A assemblies, we used recombinant baculoviruses AcR1 and AcR1Δ3’, kindly provided by Dr. Anette Schneemann (21). Upon infecting *Spodoptera frugiperda* Sf9 cells, AcR1 expresses FHV genomic RNA1 and drives robust RNA1 replication as shown by replication-dependent production of subgenomic RNA3 and its translation product protein B2 (Fig. 2A). AcR1Δ3’ expresses an FHV RNA1-derivative lacking the 3’ untranslated region required for genomic RNA1 replication and subgenomic RNA3 production. Accordingly, this RNA1 derivative translated full-length protein A but did not produce RNA replication-dependent RNA3 or its translation product B2 (Fig. 2A). Due to the high expression normally achieved with baculovirus vectors, even without RNA1 replication the protein A level in AcR1Δ3’-infected cells matched that in AcR1-infected cells. Immunofluorescence microscopy showed that most AcR1- and AcR1Δ3’-inoculated cells were baculovirus-infected and produced protein A, but dsRNA staining was only observed In cells infected by AcR1 and not AcR1Δ3’ (Fig. S1). The RNA replication defect of AcR1Δ3’ was rescued by transfecting a plasmid expressing a replicable RNA1 template, RNA1fs, that contains a frameshift mutation early in the protein A coding region and depends on independently expressed protein A for its replication (Fig. 2A). Thus, AcR1Δ3’-expressed protein A was fully capable of supporting RNA synthesis.

**Figure 2.**
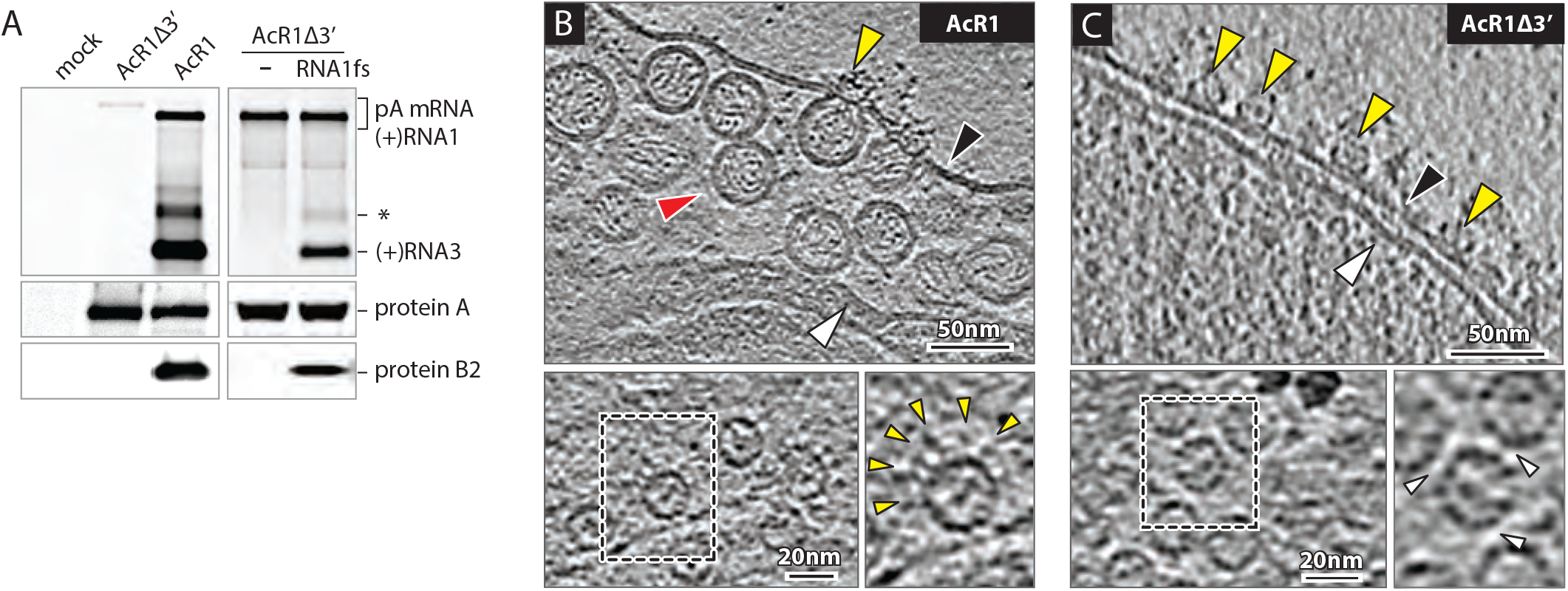
Baculovirus-launched nodavirus RNA1 replication. **(A)** Northern and western blots of Nodavirus RNAs and proteins in Sf9 cells at 48 hours post-infection with baculoviruses AcR1 or AcR1Δ3’ at an m.o.i. of 20. AcR1 launches wt RNA1 that self-replicates through cis-recognition by its product replicase protein A and produces subgenomic RNA3 and its translation product protein B2. AcR1Δ3’ lacking the RNA1 3’ noncoding region, fails to launch self-replicating RNA1, but its expressed protein A supports replication of a trans-supplied RNA1 derivative (RNA1fs) unable to produce protein A due to an N-proximal frame-shift. The FHV RNA probe also detects the primary mRNA transcript expressed from the baculovirus DNA genome and a regularly-observed virus-specific RNA species roughly twice the size of sgRNA3 (asterisk). **(B)** Cryo-ET imaging shows that AcR1-launched wt FHV RNA1 induces active RNA replication complexes bearing spherule membrane vesicles and mature crowns, while **(C)** AcR1Δ3’-expressed protein A, lacking a functional template RNA, forms crown-like structures on the OMM but no replication vesicles. Arrowheads: black = OMM, white = inner mitochondrial membrane (IMM), red = RC vesicle, yellow = crown-like structures. Higher magnification top views of the OMM show AcR1-induced crowns with leg-like extensions (yellow arrowheads), and “leg-less” and more tightly packed (white arrowheads) AcR1Δ3’ crowns.

To compare the protein A state before and during RNA replication, mitochondria from AcR1- and AcR1Δ3’-infected cells were plunge-frozen and imaged by cryo-ET. As shown in Fig, 2B, AcR1 induced mitochondrial RNA replication vesicles equivalent to those in FHV-infected *Drosophila* cells, with a high density of replication vesicles in the expanded lumen beneath the OMM, and crown-like densities at the vesicle necks. In contrast, AcR1Δ3’, expressing protein A without RNA replication, did not induce vesicles (Fig. 2C). OMM in AcR1Δ3’-infected cells bore numerous crown-like densities (Fig. 2C) not seen in mock-infected Sf9 mitochondria (Fig. S2). Top views showed that these densities were rings whose diameter and overall appearance were similar to the central turret of active 12-mer FHV replication crowns (Fig. 2B-C, lower panels). However, AcR1Δ3’-induced rings lacked the outward leg extensions of full RNA replication crowns and thereby clustered more tightly. Thus, in the absence of RNA replication, protein A forms ringed structures that are similar but not identical to FHV crowns, and without the accompanying membrane rearrangements.

Subtomogram averaging of 3001 RNA replication-active crowns on mitochondria from AcR1-infected cells produced an ~17 Å resolution structure (Fig. 3A and Fig. S3A). The resulting AcR1 crown was indistinguishable from RC crowns from FHV-infected cells (10), with a cross-correlation score of 0.977 (22). Thus, baculovirus expression recapitulates FHV RNA replication in structure and function.

**Figure 3.**
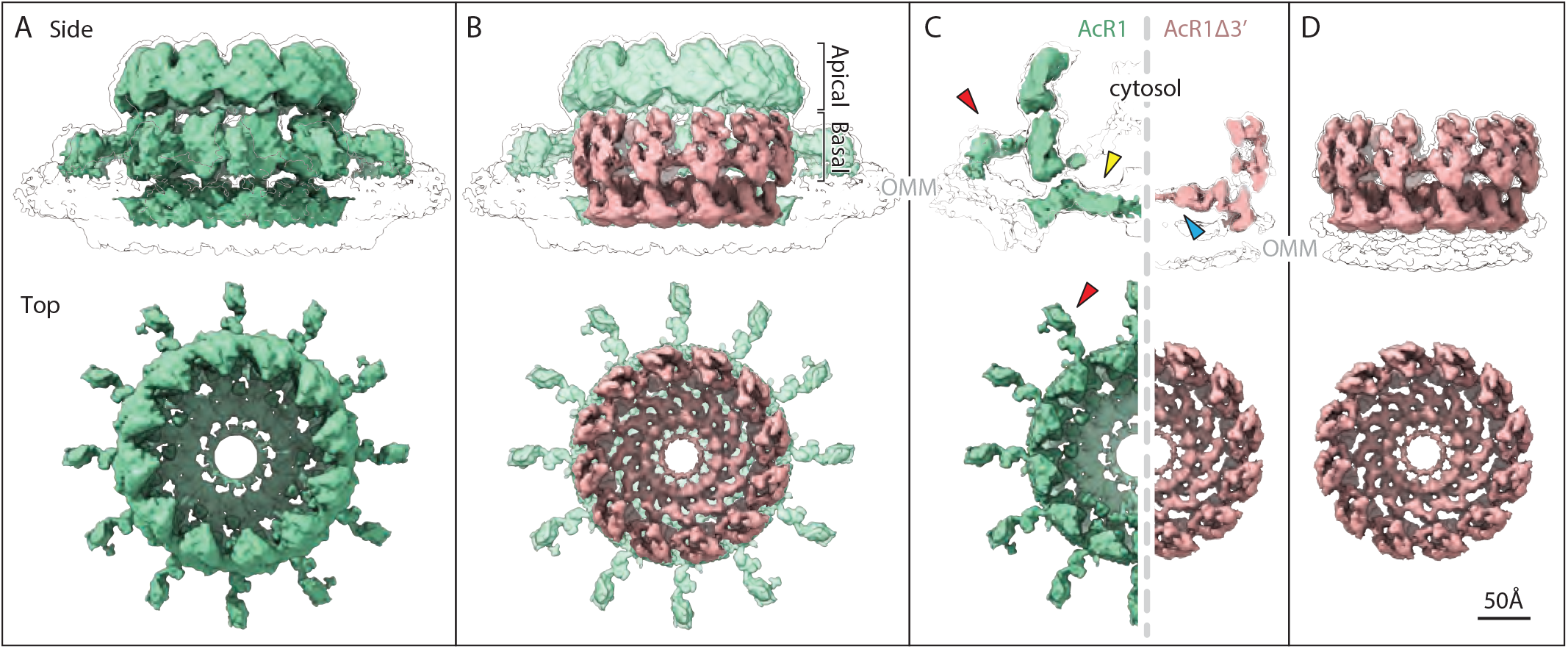
Subtomogram averages of **(A)** the baculovirus AcRI-induced, RNA replication-active mature crown (green) and **(D)** the AcR1Δ3’-induced proto-crown (pink). Upper images show side views superimposed on low-threshold density contours of the outer mitochondrial membrane (OMM), with top views below. The superposition in **(B)** and cross section views in **(C)** show how the proto-crown corresponds directly to the floor and basal lobe sections of the mature crown.

Parallel subtomogram averaging of 1685 RNA replication-inactive, crown-like particles on mitochondria from AcR1Δ3’-infected cells yielded an ~13 Å resolution structure of a short, leg-less crown-like structure with intrinsic 12-fold symmetry, hereafter designated the “proto-crown” (Fig. 3D and Fig. S3B). These pre-replication proto-crowns from AcR1Δ3’-infected Sf9 cells are markedly distinct from RNA replication-active crowns from FHV-infected *Drosophila* cells (Fig. 1D) and baculovirus AcR1-infected Sf9 cells (Fig. 3A). While preserving a 12-fold ring of subunits, the proto-crown is approximately half as tall as the mature crown and corresponds closely to the central lower portion of the mature full replication crown (Fig. 3B, movie S1). This includes the basal lobe ring and the floor, but not the outward-projecting legs and apical lobe ring of replication-active crowns, which are thus revealed as the distinct upper half of the mature crown (Fig. 3B-C). Furthermore, while the membrane under replication-active crowns curves into the semi-toroidal neck of the replication vesicle, the OMM surface below the proto-crown remains flat and unperturbed (black outlines in Fig. 3C).

### Single particle cryo-EM reveals proto-crown protein A structure

For single particle cryo-EM analysis, we used mild cross-linking (23–26) and detergent to solubilize proto-crowns from mitochondria isolated from Sf9 cells infected with a baculovirus expressing C-terminally alfa-tagged protein A (Ac-FHVptnA-alfa) (Fig. S4). Interestingly, equivalent treatment of mitochondria from S2 cells infected with a fully infectious FHV-derivative with a C-terminally alfa-tagged protein A, while bearing larger mature crowns and spherules, also yielded proto-crowns upon solubilization (Fig. S5).

2D classification showed that proto-crowns from FHV infection were 12-mer rings (Fig. S5), as expected, while proto-crowns from baculovirus-expressed protein A contained 12-mer and 11-mer rings, with C11 symmetry predominating (Fig. S4). Independent single particle analysis was conducted on C12 and C11 proto-crowns from baculovirus expression of protein A. The structures of the underlying 12- and 11-fold monomers did not reveal any significant differences. Fig. 4 presents the electron density, ribbon diagram and surface electrostatic maps of the C12 proto-crown protein A structure. Available data indicate that C12 is the native conformation of protein A on mitochondria prior to detergent solubilization. Cryo-ET shows that OMM-linked crowns and proto-crowns have C12 symmetry, while C11 assemblies have not been detected, including by 3D classification and rotational symmetry analysis ((9, 10) and this study, Fig. 3). This is functionally significant since FHV RCs on isolated mitochondria remain highly active for RNA synthesis ((6, 7) and unpublished results)). CHIKV nsP1, which is distantly related to the N-proximal half of protein A, also forms C12 rings (13, 14). Ongoing studies will examine if C11 protein A assemblies have any natural role in FHV replication or are an artifact of detergent treatment.

**Figure 4.**
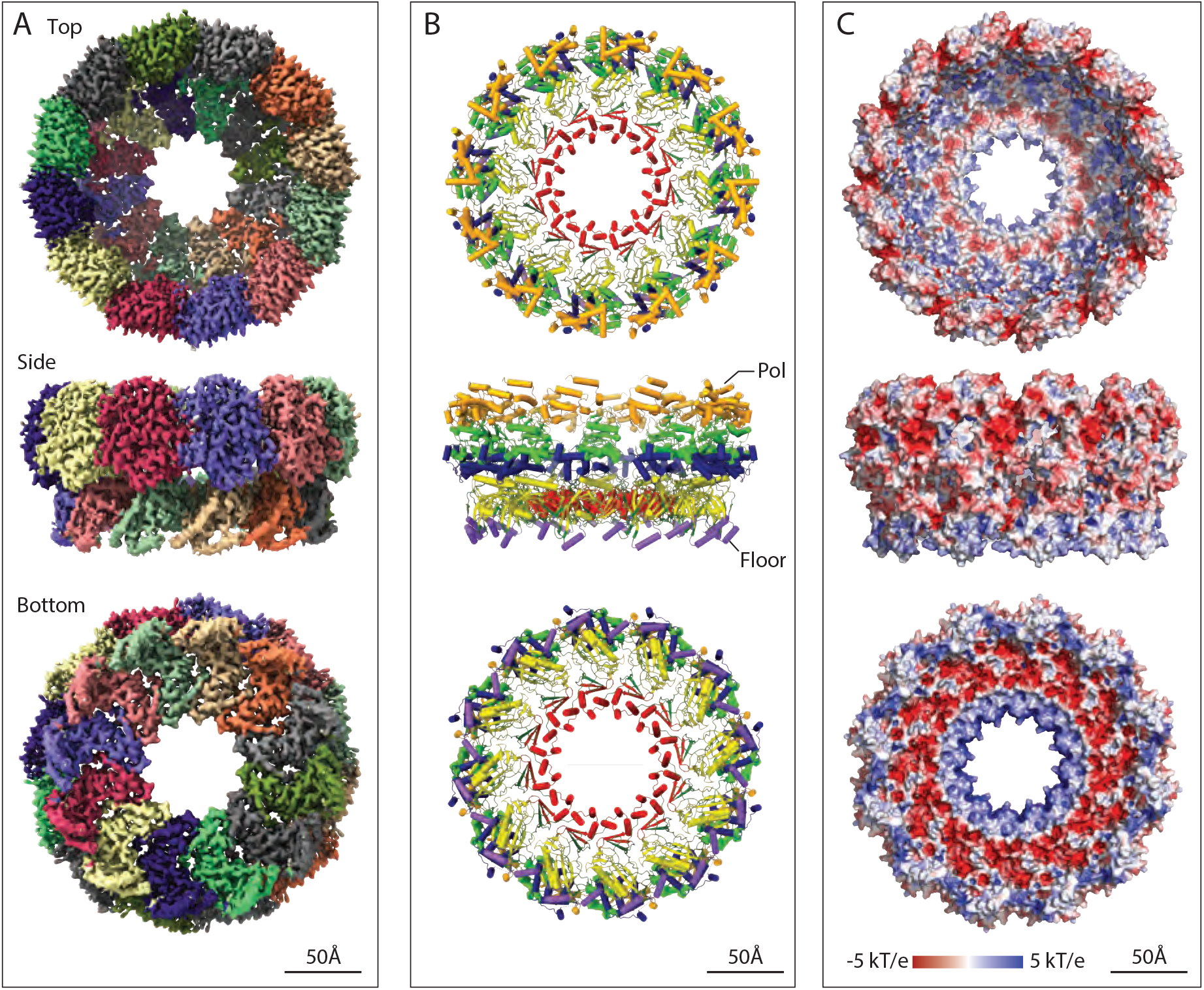
Single particle cryo-EM proto-crown structure. Top, side and bottom views of the C12 symmetric floor section in crowns from baculovirus AcR1-infected cells are shown merged with 12 copies of the Protein A Pol domain locally refined from the C12 proto-crown. **(A)** EM density maps with individually colored floor and basal lobe segments. **(B)** Ribbon diagrams of the protein A structure. The lower floor segments include domains for membrane-association (purple), capping (yellow), and central ring (red) and a functionally unassigned region (dark green). The upper Pol domain includes thumb (gold), palm (light green) and fingers (blue) domains. **(C)** Surface electrostatics calculated using APBS plugin (https://www.poissonboltzmann.org) in PyMol.

The single particle structure of the solubilized proto-crown docks precisely into the tomographic density envelopes of the mitochondrially attached proto-crown and mature crown lower central section, confirming preservation of its OMM-inserted state (Fig. 5A). As detailed further below, the repeated subunits consist of a floor domain comprising the N-proximal A_N_ segment of protein A, surmounted at its outer edge by the C-proximal protein A Pol domain (Fig. 1A and 4). The floor conformation was well maintained between proto-crowns, yielding a floor resolution of 3.1 Å (C11) to 3.2 Å (C12). This toroidal floor is nearly flat with outer and inner diameters of 19 and 7 nm and, as shown in Fig. 4B, consists from exterior to interior of three domains: a membrane interaction domain (purple), MTase/GTase RNA capping domain (yellow), and central ring with a three-strand β sheet and a bundle of three *α* helices (red). The Pol domain is more flexible (see below) but presents a typical polymerase fold.

**Figure 5.**
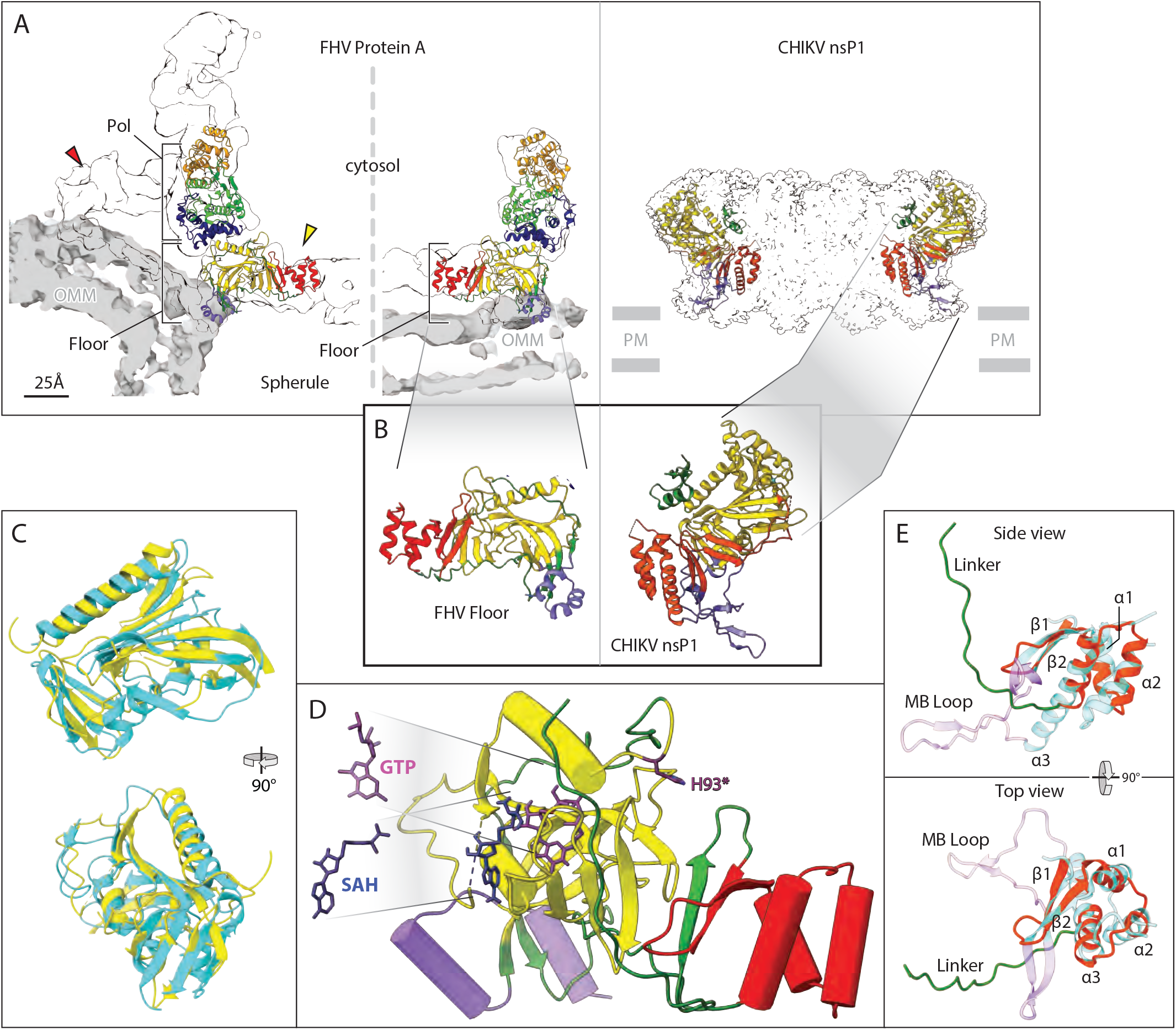
Structure of the nodavirus protein A proto-crown and selected comparisons to alphavirus nsP1. These and other zoomed-in views in figures below were taken from the highest resolution protein A structure obtained, which was the baculovirus-expressed C11 ring, but as noted in the Results no significant variations were detected between C11 and C12 subunit structures. **(A)** Fit of the atomic build of the FHV proto-crown in central cross-sections of the cryo-ET densities of the mature replication crown and the proto-crown (left side) and atomic build of the chikungunya virus (CHIKV) nsP1 crown (right side) (12). Overlaying light gray segmentations define the outer mitochondrial membrane (OMM) for the FHV crowns; positioning of the CHIKV crown on the plasma-membrane (PM) is inferred from detergent-belt in the published structure. Color coding is as defined in the legend to Figure 4B. **(B)** 12 copies of the N-terminal half of FHV protein A (segment A_N_, aa 1-452) form a flat floor in the mature crown and the proto-crown, while the equivalent CHIKV nsP1 forms a cone-shaped crown. **(C)** Superposition of the capping domains of FHV protein A (yellow) and CHIKV nsP1 (cyan). The close superposition of FHV and CHIKV capping and central channel domains and their distinct orientations in their respective crowns is further illustrated in Supplementary Movie S3. **(D)** Positions of S-Adenosyl-L-homocysteine (SAH) and GTP in the capping domain (yellow) of the FHV proto-crown floor were estimated based on superposition of the FHV protein A structural elements on the published CHIKV nsP1-ligand interacting structures and fungal pathogen E. cuniculi. The SAM/SAH binding site is composed of R100, R104, N122, and V162, while D161, Y165, P186, W215, R271 bind GTP. Note the 16 Å distance between the GTP α phosphate and GMP acceptor site H93 (dark purple). **(E)** Similarity of central ring domains in FHV protein A and CHIKV nsP1. Side- and top-views of the FHV protein A central ring domain (red) including an 18 aa linker loop (green), super-imposed on the CHIKV nsP1 RAMBO domain (cyan). Three central helical bundles (α1, α2, and α3) are sandwiched between 2 conserved upstream beta strands (β1 and β2) and the downstream polymerase linker in FHV protein A, whereas in CHIKV nsP1, the central helical bundles (α1, α2, and α3) are connected with 2 upstream beta strands (β1 and β2) followed by a membrane-binding loop (MB Loop, purple).

Overall, of the 998 aa protein A sequence, an atomic model could be built into the density from aa 55-869. C-terminal aa 897-998 are required for RNA replication but predicted to be disordered (27). Like other disordered regions, including the alphavirus nsP3 C terminus (28), this region may bind host or viral proteins or participate in other regulatory interactions (29). A few internal loops are also disordered as noted below. The following sections present the structure and mechanistic implications of the proto-crown’s functional domains in more detail.

### MTase/GTase structure and orientation in crown

Nodavirus RNAs bear 5’ m^7^Gppp caps and advanced protein sequence comparisons showed that protein A’s N-proximal region is distantly related to MTase-GTase RNA capping enzymes of the alphavirus superfamily (5). Our results confirm and extend this conclusion by showing that, despite extreme primary sequence divergence (5, 6), protein A aa 91-282 (Fig. 4B, yellow region of floor) fold into a structure highly similar to the CHIKV nsP1 MTase-GTase (13, 14), including a core six-strand β-sheet flanked by two *α*-helices and another four-strand β-sheet (Fig. 5B-C and Movie S2). Many of these structural similarities extend to the GTP N7 MTases of vaccinia virus, African swine fever virus, humans and fungal pathogen *E. cuniculi* (Movie S3).

The binding sites in protein A for GTP and the S-adenosyl homocysteine (SAH) byproduct of methyl donor S-adenosyl methionine (SAM) were inferred from superposition with the known structures of CHIKV nsP1 bound to SAH and 7-Methyl-guanosine-5’-monophosphate (m^7^GMP) (PDB: 7FGH, (30)) and *E. cuniculi* MTase bound to SAH and GTP (31) and refined by molecular dynamics simulation (Fig. 5D) (13, 14). The GTP N7 position is 5 Å from the SAH sulfur atom. However, the GTP *α* phosphate is 16 Å from the GMP acceptor site H93 (5, 6). Similar to CHIKV nsP1 (13, 14), these results imply that the protein A structure visualized corresponds to a MTase-active form while GTase function must involve significant GTP or protein movement, possibly facilitated by GTP’s position near an intersubunit interface.

While the FHV and CHIKV MTase/GTase domains share a common fold and similar GTP and SAM/SAH substrate binding sites, they differ dramatically in orientation within their respective crowns (Fig. 5A-B). The FHV proto-crown floor within which the MTase/GTase resides is essentially flat and parallel to its underlying membrane, while the corresponding CHIKV nsP1 ring is a cone expanding away from the membrane (Fig. 5A-B). As shown in Movie S2, these orientation differences correspond to a 52° rotation of the MTase-GTase domain in nsP1 relative to protein A. Whether this difference affects the exit path of 5’ capped nascent RNA from the crown remains to be determined. As noted below, the FHV and CHIKV MTase/GTase domains also differ in the position and nature of their flanking membrane-interaction domains.

### Conserved Central Ring and Diverged Membrane Interactions

The protein A sequence C-proximal to the MTase/GTase domain folds into a three-strand β-sheet followed by a bundle of three *α*-helices, which form the central ring of the proto-crown floor, bounding the 7 nm diameter channel through which RNA templates, products and nucleotides must be transported (red segment in Fig. 4B and 5 and Movie S2). The latter two β-strands and the *α*-helical bundle are conserved with CHIKV nsP1 (Fig. 5E) and conserve a few residues in similar positions. In both protein A and nsP1 the helical bundles form the central ring of the 12-mer (Fig. 5A), although their precise orientation relative to the central axis differs by a 42° rotation (Movie S2).

While the β-sheet/three helix bundle motif is conserved between protein A and nsP1, the sequences following differ significantly. The protein A sequence forms a long linker that, as discussed in the next section, runs toward the outside of the proto-crown and curves upward to eventually connect with the C-proximal polymerase domain (Fig. 5E). By contrast, in nsP1 the last *α* helix of the bundle is extended downward toward the spherule interior by 12 aa (nsp1 aa 393-404) to anchor a membrane-binding loop (Fig. 5E). This loop hooks around another membrane interaction loop inserted in the MTase/GTase domain of the adjacent nsP1, contributing to crown oligomerization and jointly forming an amphipathic unit that inserts into the membrane (13, 14).

Protein A interacts with the OMM through sequences distinct from nsP1 in position and nature, involving the N terminus (aa 1-67) and an insertion (aa 233-250) between two β-strands in the MTase/GTase. The latter insertion and N-proximal aa 55-67 each form an amphipathic helix under the outermost section of the floor. Both interact strongly with detergent in the solubilized proto-crown (Fig. S4). Similarly, modeling the protein A structure into the cryo-ET proto-crown and mature crown density maps shows both amphipathic helices interacting with the upper leaflet of the OMM (Fig. 5A, purple helices). Amino acids 1-54 were not visualized, but prior studies show that aa 15-36 are the most hydrophobic segment in protein A and act as a trans-membrane domain, localizing the protein A N-terminus to the mitochondrial intermembrane space, and that aa 1-46 are sufficient for mitochondrial targeting and membrane binding (17).

### Pol linker and floor subunit interactions

Between the three-helix bundle forming the floor central ring and the polymerase domain In protein A is an ~74 aa linker (aa 379-452). The first 18 aa (aa 379-396) are structurally well defined (Fig. 6). From the last helix of the bundle, this sequence extends under the MTase-GTase floor ~2.3 nm toward the outside of the proto-crown, then turns upward between adjacent protein subunits to near the top of the floor and base of the polymerase (Fig. 6). In this interface, this 18 aa segment provides bridging interactions to both adjoining subunits, including hydrogen bonds and electrostatic interactions. As analyzed by the PISA server (32), the interface between two adjacent floor subunits has an area of ~1804 Å^2^ and a solvation free energy of ~-21 kcal/mol. Viewed from the outside of the proto-crown (Fig. 4A-B), the interaction involves 46 residues from the left subunit and 56 residues from the right subunit, and includes 7 hydrogen bonds. Of the 46 interacting residues from the left subunit, 17 are within the polymerase linker, which thus contributes significantly to stabilizing floor assembly and might be a controlling factor in that process.

**Figure 6.**
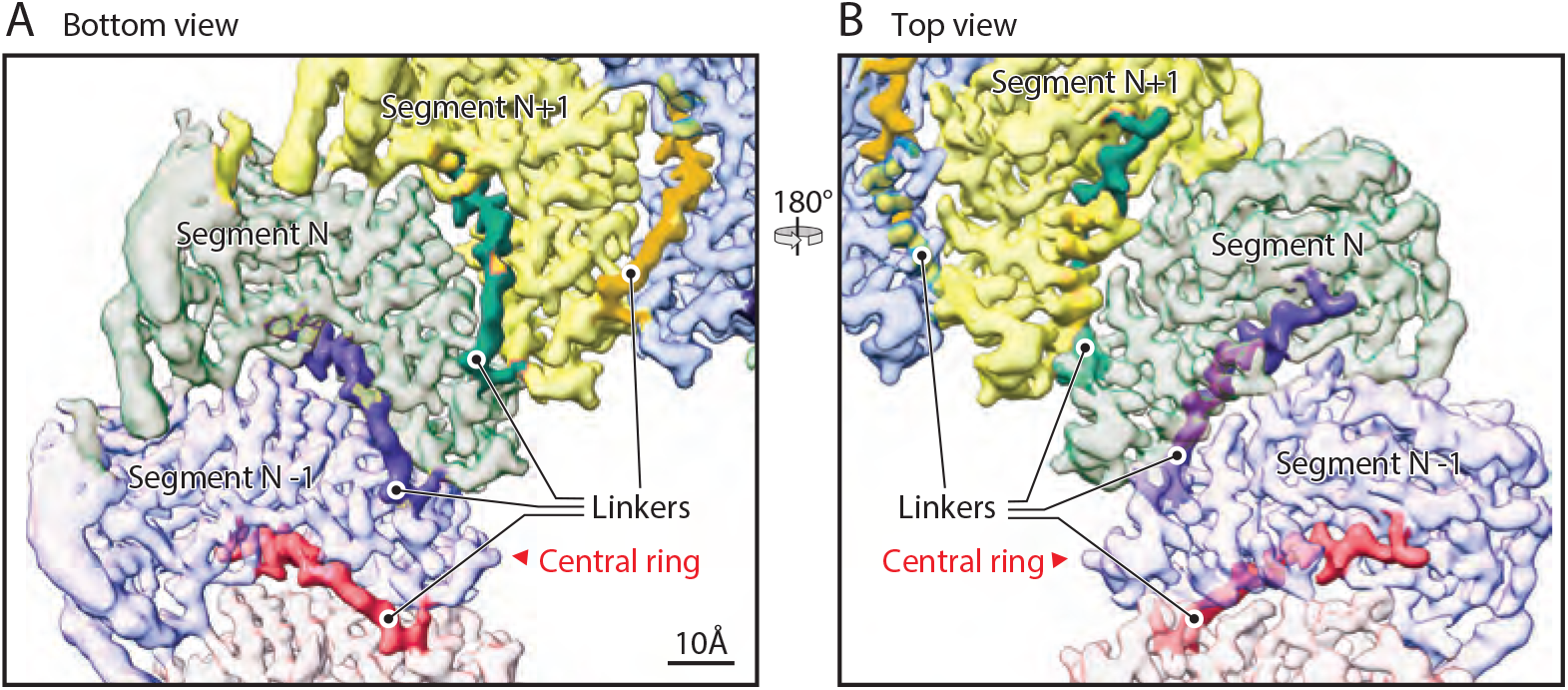
N-end of the polymerase linker is interposed between adjacent floor segments. **(A)** bottom and **(B)** top views of the crown floor density, showing the first 18 aa (dark shaded region of each colored segment) of the 74 aa linker (protein A aa 379-452) between each floor segment and its corresponding Pol domain. See main text for further details of the linker path under and between adjacent floor segments, rising to closely approach the Pol domain. Following these 18 aa, the remainder of the linker is too flexible to visualize by cryo-EM.

In contrast to the initial 18 aa’s strong participation in floor interactions, essentially all of the remaining Pol linker (aa 397-445) is too flexible to build definitively. Because of the ~27 nm potential length of this missing sequence, further studies are needed to determine if each floor domain connects with the Pol domain immediately above it as a vertical subunit, or with a Pol domain to either side in a helical array. Possible further contributions of the missing linker sequences to Pol flexibility and regulation are considered below.

### Polymerase (Pol) Structure

The Pol domain was visible from its N-terminus (aa 453), but with significantly lower resolution than the floor, becoming increasingly lower in resolution with increasing distance from the floor. Our 3D classification results indicate that the Pol domains can pivot on their contact with the floor segment below, tilting forward and backward ~5° relative to the crown’s central axis, and have inherent flexibility in their internal subdomains (Movie S4). This is similar to the highly dynamic nature of some alphavirus RNA polymerases, such as that of Ross River virus, which also possess multiple internal flexible domains (33). To improve the Pol density, we used symmetry expansion with density subtraction in *cis*TEM to isolate and individually refine the Pol structure (34, 35). We found that isolating two adjacent Pol domains for focused refinement gave the best density improvement and provided a map that, combined with the structure predicted by AlphaFold 2 (36), allowed generating a backbone trace missing the last 35 aa of the Pol domain and covering 80% of the remainder (missing aa 487-533, 630-53 and 816-28).

From its N-terminus at the bottom, adjacent to the floor, Pol folds generally upward to its C-terminus near the exposed top (Fig. 4B, 5A, 7A). Similar to other polymerases, FHV Pol has a typical “right-hand” fold with fingers, palm and thumb subdomains cupped toward the proto-crown interior (Fig. 4B, 7A). In keeping with our prior analysis (10), the DALI protein-structure comparison server (37) identified the best Pol match among known structures as Dengue flavivirus (DENV) NS5 polymerase, while the best match among alphaviruses was the Sindbis virus nsP4 polymerase (Fig. 8B-C). Thus, FHV stands as an intermediate between two important families of human and animal viruses, with an alphavirus-like RNA capping domain and flavivirus pol domain.

**Figure 7.**
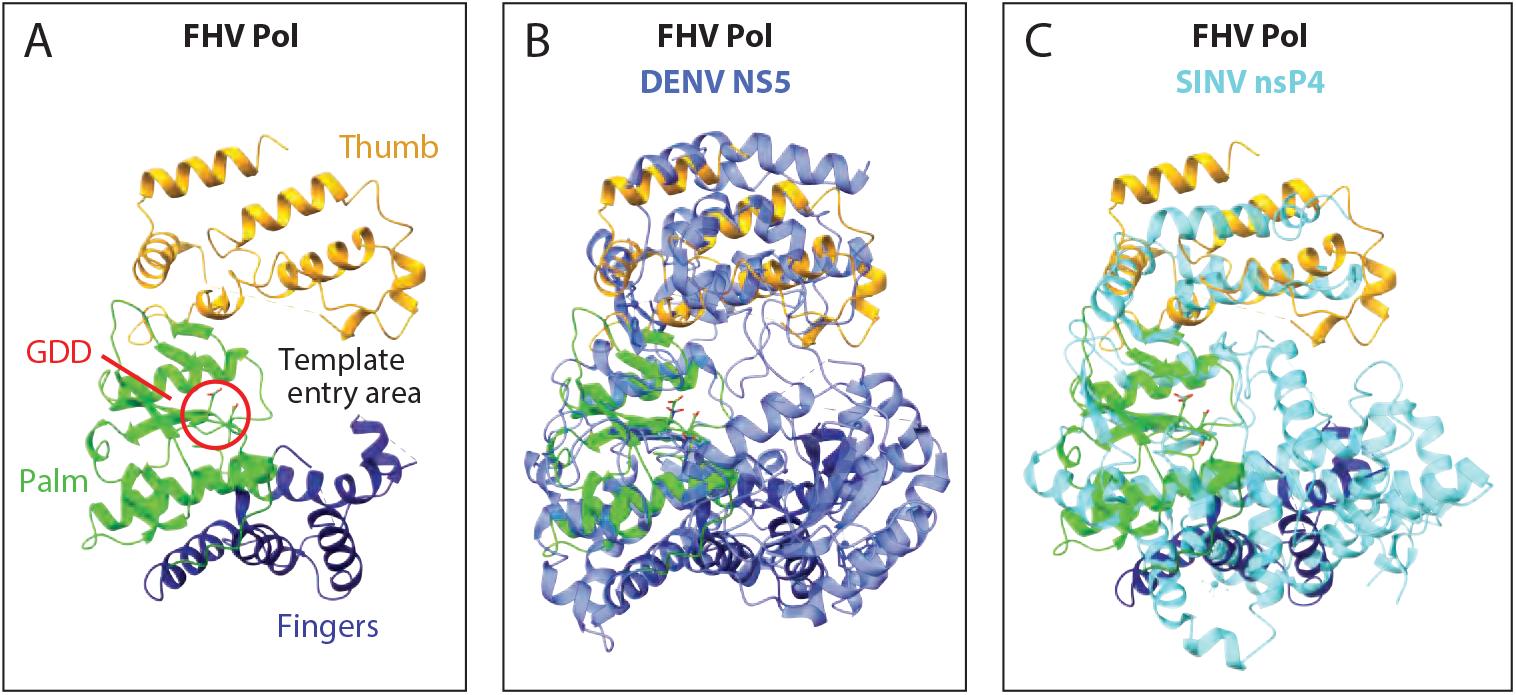
Structural comparison of Pol domains of FHV protein A, Dengue virus NS5 (PDB 4C11) and Sindbis virus nsP4 (PDB 7VW5). **(A)** Annotated ribbon diagram of the FHV protein A polymerase, in a radial view from the crown’s central axis, showing the finger (dark blue), palm (light green), and thumb regions (gold), the catalytic site (GDD, circled in red) and the RNA template entry area. **(B)** Superposition of FHV protein A Pol and Dengue virus NS5 (blue). **(C)** Superposition of FHV protein A Pol and Sindbis virus nsP4 (cyan).

**Figure 8.**
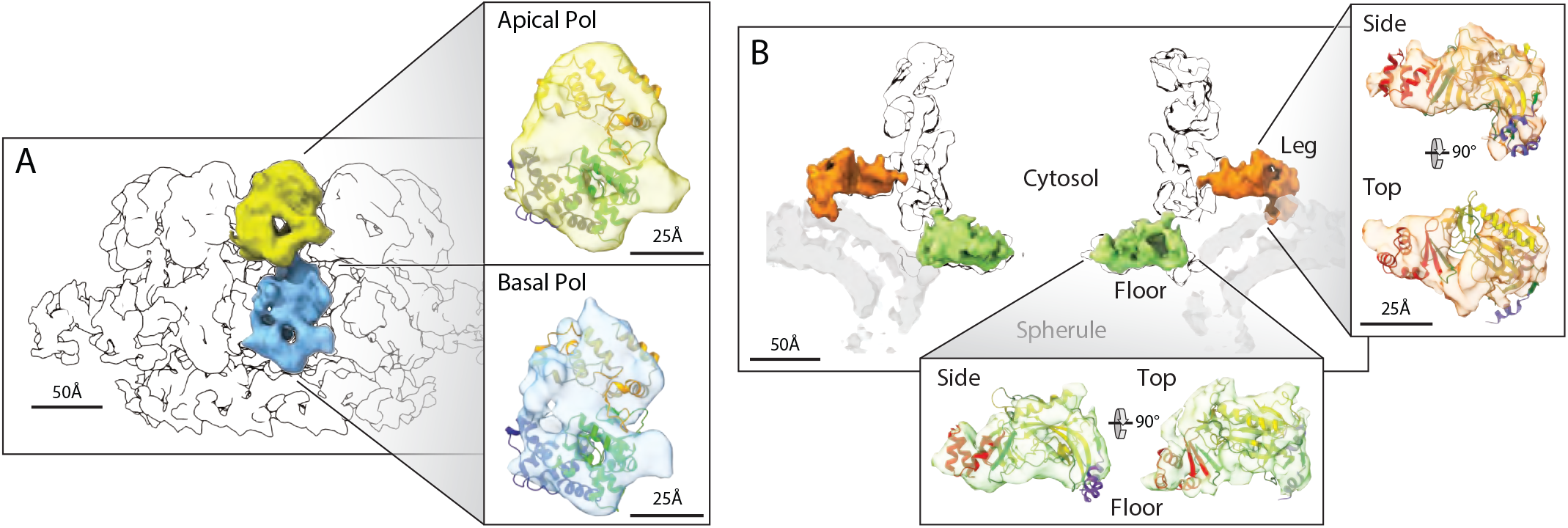
Domain assignment in the upper mature crown. **(A)** Superposition of the Pol ribbon diagram with both lobes of the central turret of the mature crown cryo-ET density, consistent with independent identification of these apical (10) and basal (Fig. 3–5) lobes as stacked Pol domains. **(B)** As shown in Fig. 3–5 and matching the A_N_ ribbon diagram superposition shown, the 12 floor segments (green) are the N-proximal A_N_ portions of protein A (aa 1-452, Fig. 1B) that are contiguous with each of the 12 basal Pol domains that constitute protein A’s C-proximal portion. Similarly, as detailed further in the main text, the legs (orange) are singled out as the A_N_ segments linked to the 12 basal Pol domains by numerous criteria, including their 12-fold repetition, position, volume, appropriately oriented membrane binding, the correspondingly good superposition of the A_N_ ribbon diagram illustrated, and the absence of any other unassigned density.

### Upper mature crown represents an alternate protein A conformation

Having found that the proto-crown constitutes the basal lobes and floor of the mature crown (Fig. 3B-C) and represents 12 copies of protein A (Fig. 4), we re-examined the remaining portions of the mature crown, i.e., the apical lobe ring and legs (Fig. 3AB). Previously we used cryo-EM visualization of a genetically inserted tag to show that the 12 apical lobes are copies of protein A’s C-proximal Pol domain (10). Thus, the full crown’s apical and basal lobe rings are stacked polymerase domains, explaining their uniform diameter. Comparing the single particle-derived Pol model to a full crown reconstruction from our prior mature FHV crown cryo-ET data (10) confirms that the the apical and basal lobe densities are highly similar, equivalently oriented in their respective rings and each fit the proto-crown Pol ribbon diagram well (Fig. 8A).

We next considered if the apical lobe Pol domains, like their basal lobe proto-crown counterparts, were linked to other portions of full-length protein A. Unlike many (+)RNA viruses, FHV does not encode a protease domain and proteolytic processing of protein A has never been reported (2). To further define the state of protein A, we performed western blotting on total protein samples from *Drosophila* S2 cells infected with the same engineered FHV derivative used in our single particle cryo-EM proto-crown studies, bearing an alfa-tag fused to protein A’s C terminus. To better examine the full protein, parallel western blots were performed with an antibody against the C-terminal alfa-tag, a monoclonal antibody against N-proximal protein A aa 99-230, and a polyclonal anti-protein A antibody that recognizes the polymerase domain (Fig. S6), presumably as well as other protein A epitopes. To identify FHV-specific bands vs. cross-reactive host bands and to estimate the relative levels of minor FHV bands, the FHV-infected cell extract was progressively diluted with uninfected cell extract.

The results confirm that >95% of protein A-derived species in infected cells are full length (112 kDa). Minor species of ~85, ~77 and ~45 kDa were visualized with the Pol-reactive polyclonal and anti-alfa-tag antibodies, but the signal for each was much less than 5% of the full-length protein A signal. Thus, since >95% of protein A in infected cells is membrane associated, predominantly in the form of crowns (10), at least 90% of the apical Pol domains in mature crowns must be linked to protein A’s N-proximal segment A_N_ (aa 1-452). However, since the apical Pol ring lacks a central floor like that formed by the N-terminal segments linked to the basal, proto-crown Pol ring, the apical A_N_ segments must be in a distinct arrangement. These upper A_N_ segments must be repeated 12 times per crown and, from the results above, should have individual volumes of ~30 nm^3^, be linked at one end to the apical polymerase base within the range of the Pol linker and at the other end linked to the OMM.

These characteristics exactly match the features of the only remaining portion of the mature crown, the legs (Fig. 8B). The 12 legs each begin just below an apical Pol domain at a distance from the estimated Pol sequence N-terminus of ~6 nm, which is close to the length of the built region of the linker (Fig. 6) and much less than the potential ~27 nm length of the full linker (aa 379-452) if unfolded (38). The estimated cryo-ET density volume of each leg is ~25 nm^3^. At its distal end, each leg interacts with the curved OMM in a fashion similar to the exterior edge of each floor segment (Fig. 5A and 8B). Moreover, rigid body fitting in ChimeraX shows that the A_N_ floor ribbon diagram from the proto-crown structure fits the cryo-ET leg density as well as the cryo-ET floor segment, and correctly positions the helical bundle start of the NTH-Pol linker at the leg’s interior end and the membrane-interacting helices at the leg’s distal membrane contact site (Fig. 8B).

These extensive matching features and the absence of any alternate, unaccounted crown density, in combination with the fact that protein A is the only viral protein required to reproduce the full structure of mature crowns from full FHV infections (11), strongly implicate the legs as the A_N_ segments that must be connected with the apical Pol segments. Due to the flexibility and potential length of the linker sequence between A_N_ and Pol, more study is needed to determine if each leg connects with the apical lobe Pol immediately above it or to one side (Fig. 8). The Fig. 8B superposition implies that the leg represents the A_N_ segment with relatively little core conformational rearrangement from the floor segment. Accordingly, relative to the floor segment, the leg is translated ~7.1 nm radially and, when viewed from outside the ring, rotated ~52° clockwise along a largely radial axis. Again, consistent with the flexibility of the Pol linker, the leg is also translated ~1.3 nm downward from the bottom of the apical Pol to interact with the adjacent basal Pol near its product RNA exit site.

### Electron density in the crown central channel

As we noted previously, the Pol domains in the central turret (Fig. 8A) appear inconsistent with (+)RNA synthesis, due to their physical separation from the dsRNA template beneath the crown floor and other issues. Rather, the requirements of (+)RNA synthesis appear to demand an alternate state of protein A near the 7 nm central channel of the crown floor to access the viral dsRNA template inside the spherule (1, 10). In keeping with this, strong electron density spanning the central channel of the mature crown floor has been consistently observed in all cryo-EM studies of replication-active nodavirus crowns ((9–11) and this paper). To better examine this density, we locally refined the inner central ring region of the floor of cryo-ET-imaged crowns from FHV-infected cells without imposed symmetry. Fig. 9 shows this refined central density, and its asymmetric contacts with the inner ring of the floor, within the sub-tomogram average of the full, mature crown also with no imposed symmetry. The estimated 41 nm^3^ volume of this central density appears reasonably consistent with the ~44 nm^3^ volume of individual Pol domains in the basal and apical rings, particularly considering their relatively low resolution. Possible origins of this density are considered further in the Discussion.

**Figure 9.**
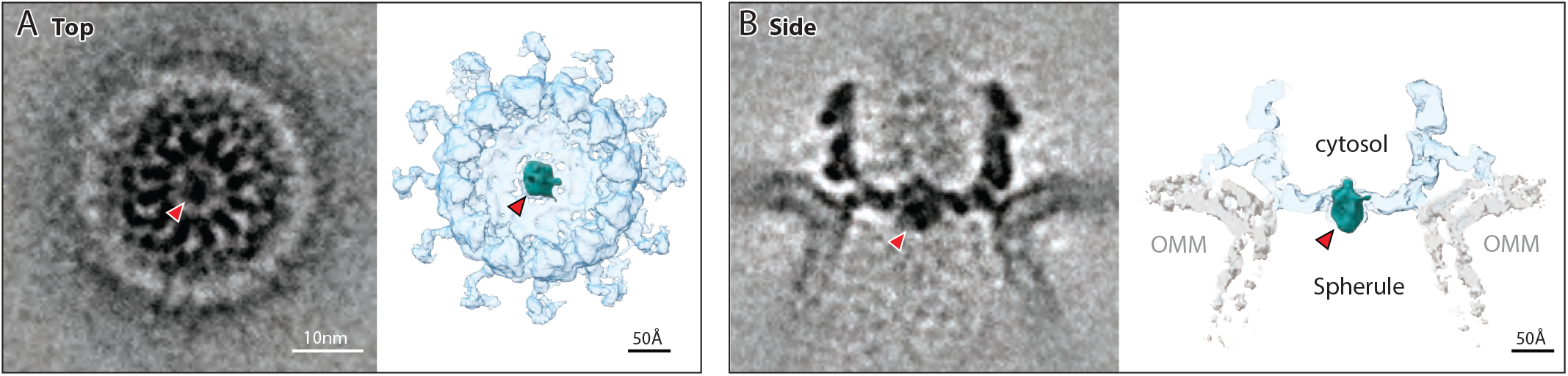
Asymmetric density in the crown floor central channel. **(A)** Top and **(B)** side views of the cryo-ET density of the RNA replication-active mature crown from FHV-infected cells refined by sub-tomogram averaging without imposed symmetry. In both graded density (left) and densi-ty-thresholded views (right), red arrowheads point to an asymmetric density within the 7 nm diameter crown floor central channel, distinct from the floor segment densities of Figs. 3–6. This density (colored teal in the right side views) occupies a volume closely similar to that of the apical and basal polymerase lobes.

## Discussion

The results presented here show that the full, mature nodavirus RNA replication crown is a 24-mer of protein A in two distinct conformations. Twelve copies of protein A constitute a lower, proto-crown precursor that forms on the OMM when protein A is expressed alone, in the absence of a replicable RNA template (Fig. 3 and 4). Active RCs functional for repeated (+)RNA synthesis, the major job of RNA replication, always possess a second 12-mer ring of protein A in a distinct conformation above and surrounding the proto-crown (Fig. 3 and 8), as well as an invaginated spherule vesicle containing a viral dsRNA template (Fig. 2B).

Forming nodavirus RNA replication vesicles requires protein A, a functional RNA template and RNA synthesis (20), suggesting that the vesicle is generated by being filled with the dsRNA product of (-)RNA synthesis (9, 12). The resulting replication vesicle, pressurized by negatively charged dsRNA, is held together at the neck by the mature crown, which depends on its dual rings of membrane interaction sites in the upper crown legs as well as the lower proto-crown floor to closely constrain and stabilize the curved membrane neck (Fig. 3A and (10)). Thus, (-)RNA synthesis, invaginating the spherule replication vesicle and maturing the pre-replication proto-crown to the full double ring crown are closely linked events, and possibly also closely linked to recruiting the founding (+)RNA template for (-)RNA synthesis. Determining the precise order and interrelation of these steps will be highly informative. Below we discuss further aspects of the nodavirus crown structure, mechanistic implications of these features for RNA replication, and broader implications for (+)RNA viruses.

### Extended Pol linker and central channel electron density

A surprising feature of the protein A structure is the 74 aa sequence (aa 379-452) linking the floor’s central ring to Pol. In the proto-crown, the initial 18 aa of this sequence (aa 379-396) form a well-defined elbow that reaches from the three-helix bundle of the central ring to near the base of Pol and contributes significantly to the intersubunit interactions that assemble the proto-crown floor (Fig. 6). Re-positioning this connecting sequence when segment A_N_ shifts relative to Pol between its multimerized inner floor assembly and separated outer leg positions (Fig. 8) might be coupled to dissolving the floor interactions.

Although this elbow reaches within ~2 nm of the Pol start domain in the proto-crown structure, there remain 56 aa (aa 397-452) of primary sequence between the elbow’s end and Pol start. This additional linker sequence was too flexible to resolve in the structure and, if extended, could stretch over 22 nm, greatly exceeding the ~7 nm movement of A_N_ relative to Pol between its proto-crown floor and upper crown leg positions (Fig. 8). In principle, this flexible linker might support Pol movement anywhere within the crown for (+)RNA synthesis, (-)RNA synthesis or other steps.

Fig. 9 shows that in mature nodavirus crowns active in (+)RNA synthesis, the crown floor’s central channel is occupied by an asymmetric electron density with a volume similar to the protein A Pol domain. This central density has been visualized in all previous cryo-ET reconstructions of mature crowns from FHV-infected cells (9–11), and was present in all 3D classes of mature crowns obtained in the current analysis. This is of considerable interest since the 24 Pol domains of the central turret (Fig. 8A) are blocked by the crown floor from accessing the spherule-protected dsRNA template, and therefore incompatible with (+)RNA synthesis. These considerations implied that crowns must contain another form of Pol with direct access to the dsRNA (1, 10, 12), which a Pol in the central channel would provide. As summarized in the next section, emerging results on CHIKV crowns strongly support this hypothesis.

A Pol domain might be recruited to the central channel in at least two ways. First, a single Pol domain in the apical or basal ring might disengage and, flexibly tethered by the long Pol linker, bend down to engage the central channel. Consistent with this, Pol variability in our 3D classification (see Results and Movie S4) shows that these domains are not strongly bound in the central turret. Initial efforts to search for crowns that might be missing one or more apical or basal Pol domains have been inconclusive to date, and efforts in this area are ongoing. The central channel density might alternatively represent a 25^th^ Pol in the crown. Such a 25^th^ Pol might be a rare independent Pol processed away from other protein A segments (Fig. S6) or part of a full-length protein A. A 25^th^ full length protein A might have the A_N_ membrane interaction domains neutralized by presently unknown interactions or utilize the long, flexible linker between A_N_ and Pol to interact with the spherule membrane below the crown floor.

Beyond (+)RNA synthesis and in addition to their structural roles, the Pol domains in the proto-crown or mature crown central turrets likely function in other required Pol functions, including RNA template recruitment and (-)RNA synthesis (2, 19, 20), and possibly bridging interactions to downstream processes like translation and encapsidation (see next section).

### Relation to alphavirus RNA replication complexes

While this report was in final preparation, a manuscript uploaded to *bioRxiv* presented further results on CHIKV RNA replication protein structure, interaction and relation to cryo-ET images of *in vivo* RCs (39). This included the single particle cryo-EM structure of an nsP1+nsP2+nsP4 complex, comprising one copy of nsP4 polymerase bound to the central pore of the nsP1 12-mer ring (Fig. 5A), with one nsP2 helicase domain bound to the cytosolic side of nsP4. This nsP1+nsP2+nsP4 complex fit well into the membrane-proximal ~7 nm portion of ~17 nm tall cryo-ET-imaged crowns at the necks of spherule RNA RCs in CHIKV-infected cells (39). The composition of the upper, cytosol-facing, 12-fold symmetric ~10 nm of these crowns was not defined but was suggested to include CHIKV RNA replication factor nsP3 and possibly host factors. In addition to 17 nm tall crowns on full RCs, late-stage infected cells contained shorter crown-like rings that appeared to correspond to the nsP1+nsP2+nsP4 complex alone, which were always found on flat membranes lacking spherule invaginations and their dsRNA contents.

These alphavirus findings show striking parallels with the nodavirus results reported here, including multi-stage assembly of the RNA replication crown by stacking distinct 12-fold symmetric protein rings. Prior to RNA synthesis, both viruses first assemble single ring precursors on flat membranes, and then mature these into active RCs by invaginating membrane spherules filled with dsRNA and adding a further, membrane-distal protein layer to the crown precursor.

Nodaviruses and alphaviruses both require viral RNA synthesis for membrane spherule formation (9, 20, 40), and current and prior nodavirus results further document that the crown assembly transition revealed here from proto-crown precursor to replication-active mature crown is triggered by RNA template availability (Fig. 2–3). Potential broad significance for this RNA-activated assembly transition is supported by parallels with an RNA-dependent RNA polymerase extract from brome mosaic virus (BMV), an alphavirus superfamily member with RNA replication proteins related to CHIKV nsP1, 2 and 4. This detergent-solubilized BMV polymerase complex lacks functional endogenous RNA templates and requires *in vitro*-supplied viral RNA templates. Nevertheless, template-dependent polymerase activity can only be isolated from cells expressing viral RNA replication proteins plus a viral RNA template; cells lacking an RNA template yield extracts with the same complement of viral proteins but no activity on added templates (41). Thus, without a functional template, replication proteins fail to complete assembly to an active state.

Conservation of multiple key principles of RC structure, assembly and function between nodaviruses and alphaviruses is particularly notable given the considerable evolutionary distance between their RNA replication proteins, which are dramatically diverged in size, complexity, expression strategy and sequence (Fig. 1A and (5)). Preservation of these common features thus implies that the underlying RNA replication principles are ancient and likely to extend to other (+)RNA viruses. Flaviviruses, e.g., also form spherule RCs (42, 43) that might embody at least some of these features, particularly since the protein A structure reveals FHV as a missing link encoding an alphavirus-like MTase-GTase and a flavivirus-like polymerase (Fig. 5 and 7). Moreover, despite even greater evolutionary differences, coronaviruses also form membrane-spanning multimeric crowns that channel release of progeny RNAs to the cytosol (1, 10, 15, 16). The crown and RC features revealed here also underscore and further illuminate multiple parallels with the replicative cores of dsRNA virus and retrovirus virions (1).

Complementing the conserved noda- and alphavirus features are differences revealing spherule operational flexibilities. While conserving their core structure and 12-mer assembly, e.g., the MTase-GTase and central ring domains differ in structure and attachment points of their membrane interaction domains (Fig. 5) and assemble into a flat floor for protein A and a cone in nsP1 (Fig. 5A-B). Also intriguing is the absence in nodaviruses of the NTPase/helicase domain conserved across the alphavirus-like superfamily (Fig. 1), where it is required for viral template RNA recognition and recruitment, ongoing synthesis of (+), (-) and subgenomic RNAs, and 5’ RNA triphosphatase activity in RNA capping (44–46). To provide corresponding functions, nodaviruses might recruit one or more host proteins. Since nearly all cryo-ET crown density is assigned to protein A (Fig. 8), any such host protein(s) presumably would be in low copy number relative to protein A, and might contribute partially to the asymmetric density in the floor central channel (Fig; 8). Alternatively, some helicase functions might be replaced by polymerase dsRNA unwinding activity (47) or unconventional activities of other protein A domains. Overall, the ability of the single, 998 aa FHV protein A to provide varied structural and functional roles paralleling those of the 2474 aa of CHIKV nsP1+2+3+4 within multi-layered crowns is a telling example of viral protein multifunctionality and plasticity. Further studies should illuminate the elegant adaptations, including protein A’s alternate conformational modes and likely its flexible Pol linker, that allow nodaviruses to utilize their unusually small and simple genomes with great evolutionary success.

Among many novel questions posed by the results presented here are the functions of the mature crown upper ring. Possibilities include shielding RNA replication processes, templates and intermediates from innate immune recognition, assisting transfer of RNA products into translation and encapsidation, and providing a reaction chamber to facilitate RNA capping by regulating the rate and path of nascent product release. The proto-crown and crown structures reported here provide important foundations for addressing these and other crucial questions that should greatly enhance understanding and control of (+)RNA viruses.

## Materials & Methods

Detailed descriptions of cell culture, viruses, infections, transfections, RNA and protein analysis, immunofluorescence and cryo-EM are provided as Supporting Information.

## Data availability

All data are included in the article and/or supporting information. The coordinates of the atomic models of the FHV protein A floor segment refined using C11 and C12 cryoEM density maps and of the atomic model of the FHV protein A Pol domain are being deposited in the Protein Data Bank (PDB). The cryo-EM maps of the C11 and C12 FHV protein A floor segment and the associated locally refined Pol density map are being deposited in the EM Data Bank (EMDB). Accession codes will be provided at the time of final publication.

## Acknowledgments

We thank the UW-Madison Cryo-EM Research Center and OHSU Pacific Northwest Cryo-EM Center for exceptional facilities and data collection support, the Morgridge Research Computing group for outstanding computational support, Anette Schneemann for recombinant baculoviruses, Adam Steinberg for creative graphics assistance, members of our groups for valuable discussions, and Jean-Yves Sgro for animation assistance. Some of this work was performed in the Cryo-EM Research Center (CEMRC) in the Department of Biochemistry at the University of Wisconsin-Madison. A portion of this research was supported by NIH grant U24GM129547 and performed at the Pacific Northwest Cryo-EM Center at Oregon Health and Science University and accessed through EMSL (grid.436923.9), a DOE Office of Science User Facility sponsored by the Office of Biological and Environmental Research. This research was performed using the compute resources and assistance of the UW-Madison Center For High Throughput Computing (CHTC) in the Department of Computer Sciences. The CHTC is supported by UW-Madison, the Advanced Computing Initiative, the Wisconsin Alumni Research Foundation, the Wisconsin Institutes for Discovery, and the National Science Foundation, and is an active member of the Open Science Grid, which is supported by the National Science Foundation and the U.S. Department of Energy’s Office of Science. P.A. and T.G. are investigators of the Morgridge Institute for Research and the director and an investigator, respectively, of the John and Jeanne Rowe Center for Research in Virology, and gratefully acknowledge their support.

## References

1. M. Nishikiori, J. A. den Boon, N. Unchwaniwala, P. Ahlquist, Crowning Touches in Positive-Strand RNA Virus Genome Replication Complex Structure and Function. Annu. Rev. Virol. (2022) https://doi.org/10.1146/annurev-virology-092920-021307.

2. P. A. Venter, A. Schneemann, Recent insights into the biology and biomedical applications of Flock House virus. Cell. Mol. Life Sci. CMLS 65, 2675–2687 (2008).

3. B. D. Price, R. R. Rueckert, P. Ahlquist, Complete replication of an animal virus and maintenance of expression vectors derived from it in Saccharomyces cerevisiae. Proc. Natl. Acad. Sci. U. S. A. 93, 9465–9470 (1996).

4. R. Lu, et al., Animal virus replication and RNAi-mediated antiviral silencing in Caenorhabditis elegans. Nature 436, 1040–1043 (2005).

5. T. Ahola, D. G. Karlin, Sequence analysis reveals a conserved extension in the capping enzyme of the alphavirus supergroup, and a homologous domain in nodaviruses. Biol. Direct 10, 16 (2015).

6. T. Quirin, Y. Chen, M. K. Pietilä, D. Guo, T. Ahola, The RNA Capping Enzyme Domain in Protein A is Essential for Flock House Virus Replication. Viruses 10, E483 (2018).

7. B. G. Kopek, G. Perkins, D. J. Miller, M. H. Ellisman, P. Ahlquist, Three-dimensional analysis of a viral RNA replication complex reveals a virus-induced mini-organelle. PLoS Biol. 5, e220 (2007).

8. D. J. Miller, M. D. Schwartz, P. Ahlquist, Flock house virus RNA replicates on outer mitochondrial membranes in Drosophila cells. J. Virol. 75, 11664–11676 (2001).

9. K. J. Ertel, et al., Cryo-electron tomography reveals novel features of a viral RNA replication compartment. eLife 6, e25940 (2017).

10. N. Unchwaniwala, et al., Subdomain cryo-EM structure of nodaviral replication protein A crown complex provides mechanistic insights into RNA genome replication. Proc. Natl. Acad. Sci. U. S. A. 117, 18680–18691 (2020).

11. J. A. den Boon, et al., Multifunctional Protein A is the Only Viral Protein Required for Nodavirus RNA Replication Crown Formation. Viruses 14, 2711 (2022).

12. N. Unchwaniwala, H. Zhan, J. A. den Boon, P. Ahlquist, Cryo-electron microscopy of nodavirus RNA replication organelles illuminates positive-strand RNA virus genome replication. Curr. Opin. Virol. 51, 74–79 (2021).

13. R. Jones, G. Bragagnolo, R. Arranz, J. Reguera, Capping pores of alphavirus nsP1 gate membranous viral replication factories. Nature 589, 615–619 (2021).

14. K. Zhang, et al., Structural insights into viral RNA capping and plasma membrane targeting by Chikungunya virus nonstructural protein 1. Cell Host Microbe 29, 757–764.e3 (2021).

15. G. Wolff, et al., A molecular pore spans the double membrane of the coronavirus replication organelle. Science 369, 1395–1398 (2020).

16. N. Unchwaniwala, P. Ahlquist, Coronavirus dons a new crown. Science 369, 1306–1307 (2020).

17. D. J. Miller, P. Ahlquist, Flock house virus RNA polymerase is a transmembrane protein with aminoterminal sequences sufficient for mitochondrial localization and membrane insertion. J. Virol. 76, 9856–9867 (2002).

18. P. M. Van Wynsberghe, P. Ahlquist, 5’ cis elements direct nodavirus RNA1 recruitment to mitochondrial sites of replication complex formation. J. Virol. 83, 2976–2988 (2009).

19. P. M. Van Wynsberghe, H.-R. Chen, P. Ahlquist, Nodavirus RNA replication protein a induces membrane association of genomic RNA. J. Virol. 81, 4633–4644 (2007).

20. B. G. Kopek, E. W. Settles, P. D. Friesen, P. Ahlquist, Nodavirus-induced membrane rearrangement in replication complex assembly requires replicase protein a, RNA templates, and polymerase activity. J. Virol. 84, 12492–12503 (2010).

21. P. A. Venter, N. K. Krishna, A. Schneemann, Capsid protein synthesis from replicating RNA directs specific packaging of the genome of a multipartite, positive-strand RNA virus. J. Virol. 79, 6239–6248 (2005).

22. E. F. Pettersen, et al., UCSF ChimeraX: Structure visualization for researchers, educators, and developers. Protein Sci. Publ. Protein Soc. 30, 70–82 (2021).

23. V. Q. Nguyen, et al., Molecular architecture of the ATP-dependent chromatin-remodeling complex SWR1. Cell 154, 1220–1231 (2013).

24. J. Y. Kang, et al., Structural basis of transcription arrest by coliphage HK022 Nun in an Escherichia coli RNA polymerase elongation complex. eLife 6, e25478 (2017).

25. D. Y. Zhao, et al., Cryo-EM structure of the native rhodopsin dimer in nanodiscs. J. Biol. Chem. 294, 14215–14230 (2019).

26. Y. Xie, et al., Cryo-EM structure of the yeast TREX complex and coordination with the SR-like protein Gbp2. eLife 10 (2021).

27. L. Gitlin, T. Hagai, A. LaBarbera, M. Solovey, R. Andino, Rapid evolution of virus sequences in intrinsically disordered protein regions. PLoS Pathog. 10, e1004529 (2014).

28. V. Kril, O. Aïqui-Reboul-Paviet, L. Briant, A. Amara, New Insights into Chikungunya Virus Infection and Pathogenesis. Annu. Rev. Virol. 8, 327–347 (2021).

29. P. E. Wright, H. J. Dyson, Intrinsically disordered proteins in cellular signalling and regulation. Nat. Rev. Mol. Cell Biol. 16, 18–29 (2015).

30. K. Zhang, et al., Molecular basis of specific viral RNA recognition and 5’-end capping by the Chikungunya virus nsP1. Cell Rep. 40, 111133 (2022).

31. C. Fabrega, S. Hausmann, V. Shen, S. Shuman, C. D. Lima, Structure and mechanism of mRNA cap (guanine-N7) methyltransferase. Mol. Cell 13, 77–89 (2004).

32. E. Krissinel, K. Henrick, Inference of macromolecular assemblies from crystalline state. J. Mol. Biol. 372, 774–797 (2007).

33. Y. B. Tan, et al., Crystal structures of alphavirus nonstructural protein 4 (nsP4) reveal an intrinsically dynamic RNA-dependent RNA polymerase fold. Nucleic Acids Res. 50, 1000–1016 (2022).

34. J. Jung, et al., High-resolution cryo-EM structures of outbreak strain human norovirus shells reveal size variations. Proc. Natl. Acad. Sci. U. S. A. 116, 12828–12832 (2019).

35. T. Grant, A. Rohou, N. Grigorieff, cisTEM, user-friendly software for single-particle image processing. eLife 7 (2018).

36. J. Jumper, et al., Highly accurate protein structure prediction with AlphaFold. Nature 596, 583–589 (2021).

37. L. Holm, L. M. Laakso, Dali server update. Nucleic Acids Res. 44, W351–355 (2016).

38. S. R. K. Ainavarapu, et al., Contour length and refolding rate of a small protein controlled by engineered disulfide bonds. Biophys. J. 92, 225–233 (2007).

39. Y. B. Tan, et al., Molecular Architecture of the Chikungunya Virus Replication Complex. bioRxiv, 2022.04.08.487651 (2022).

40. K. Kallio, et al., Template RNA length determines the size of replication complex spherules for Semliki Forest virus. J. Virol. 87, 9125–9134 (2013).

41. R. Quadt, M. Ishikawa, M. Janda, P. Ahlquist, Formation of brome mosaic virus RNA-dependent RNA polymerase in yeast requires coexpression of viral proteins and viral RNA. Proc. Natl. Acad. Sci. U. S. A. 92, 4892–4896 (1995).

42. S. Welsch, et al., Composition and three-dimensional architecture of the dengue virus replication and assembly sites. Cell Host Microbe 5, 365–375 (2009).

43. L. K. Gillespie, A. Hoenen, G. Morgan, J. M. Mackenzie, The endoplasmic reticulum provides the membrane platform for biogenesis of the flavivirus replication complex. J. Virol. 84, 10438–10447 (2010).

44. X. Wang, et al., Brome mosaic virus 1a nucleoside triphosphatase/helicase domain plays crucial roles in recruiting RNA replication templates. J. Virol. 79, 13747–13758 (2005).

45. P. A. Kroner, B. M. Young, P. Ahlquist, Analysis of the role of brome mosaic virus 1a protein domains in RNA replication, using linker insertion mutagenesis. J. Virol. 64, 6110–6120 (1990).

46. L. Vasiljeva, A. Merits, P. Auvinen, L. Kääriäinen, Identification of a novel function of the alphavirus capping apparatus. RNA 5’-triphosphatase activity of Nsp2. J. Biol. Chem. 275, 17281–17287 (2000).

47. M. W. Cho, O. C. Richards, T. M. Dmitrieva, V. Agol, E. Ehrenfeld, RNA duplex unwinding activity of poliovirus RNA-dependent RNA polymerase 3Dpol. J. Virol. 67, 3010–3018 (1993).

