## Supplemental Information for "Nodavirus RNA Replication Crown Architecture Reveals Proto-Crown Precursor and Viral Protein A Conformational Switching"

**This PDF file includes:**

Supporting Information Text (Methods)
Figures S1 to S6
Tables S1and S2
SI References

### Supporting Information Text

#### Methods

**Cell culture.** *Drosophila melanogaster* S2 cells were maintained at 28°C in ExpressFive serum-free medium containing 120 U/ml penicillin, 100 µg/ml streptomycin, and 2mM L-glutamine (Pen/Strep/Glu). *Spodoptera frugiperda* cell line Sf9 was grown in Sf-900 III media either stationary or shaking at 120 rpm at 28°C.

**Infections.** S2 cells were infected with FHV at m.o.i. of 10 in Schneider's medium containing Pen/Strep/Glu supplemented with 10% bovine calf serum (BCS) and incubated for 16 hours at 28°C prior to analysis. Infections of Sf9 cells with recombinant baculoviruses AcR1 and AcR1Δ3' (kind gifts from Anette Schneemann, originally named AcR1δ and AcR1-3'UTR, respectively in (1)) or Ac-FHVptnAalfa (newly generated, see below) were performed at m.o.i. of 20 by gently rocking on an end-to-end shaker for 1 hour at room temperature prior to transferring the cells to a shaking culture flask and incubating for 48-72 hours at 28°C.

**Recombinant DNA and DNA transfections.** Recombinant FHV-Aalfa was generated via co-transfection of S2 cells with pIE1hr5-FHVRNA1Rz-ptnAalfa-noB1 to launch a replicable genomic RNA1 expressing C-terminally alfa-tagged protein A and a second plasmid pMT-FHVRNA2Rz to launch a wildtype replicable genomic RNA2, using MIRUS TransIT transfection reagents. Recombinant baculovirus Ac-FHVptnAalfa expressing C-terminally alfa-tagged protein A was generated through co-transfection of Sf9 cells with baculovirus transfer plasmid pOET-FHVptnAalfa and a baculovirus "genomic backbone" using MIRUS transfection and *flash*Bac technology according to instructions. Plasmids were constructed using standard molecular cloning technology using restriction and ligation enzyme-based procedures including PCR and synthetic DNA-based methods to generate engineered DNA fragments. Oligonucleotides and gBLOCKs were purchased from IDT DNA Technologies. Plasmid sequences were verified using thermocycling-based BigDye sequencing technology and analysis at the University of Wisconsin-Madison Biotechnology Center Core Sequencing Facilities.

**Protein extraction and Western blotting.** Total protein was extracted using preheated (85°C) Cracking Buffer (8M Urea, 5% SDS, 40 mM TrisHCl pH 7.0, 100 mM EDTA, 25% glycerol, and a trace of orange G) at 200 µl per ~1 x $10^6$ cells. Typically, 10-15 µl sample re-heated to 85°C for 10 minutes prior to loading was resolved in NuPAGE 4-12% or 8% Bis-Tris Midi Protein denaturing SDS PAGE gels (Thermo Fisher Scientific) using accompanying NuPAGE SDS running buffer conditions. Upon completion, gels were washed in $ddH_2O$ for 10 minutes and equilibrated in 2x NuPAGE transfer buffer for 15 minutes. Protein was transferred to Immobilon-FL PVDF membrane pre-wetted in methanol and equilibrated in 2x NuPAGE transfer buffer using semi-dry blotting for 10 minutes at 25V. Membranes were allowed to dry and rehydrated using methanol, $ddH_2O$, and PBS washes prior to incubation in Odyssey Blocking Buffer for 1 hour at room temperature. Blocked membranes were incubated with primary antibody, typically at 1:1000 to 1:5000 dilution, in Odyssey Blocking Buffer supplemented with 0.2% Tween20 for 1-4 hours at room temperature. Membranes were washed three times in Odyssey Blocking Buffer/0.2% Tween20 for 5 minutes at room temperature and incubated with secondary antibody conjugated to IRDye 680RD or 800CW (1:10,000) in Odyssey Blocking Buffer supplemented with 0.2% Tween20 and 0.01 % SDS for 1 hour at room temperature. Membranes were again washed three times in Odyssey Blocking Buffer/0.2% Tween20 for 5 minutes at room temperature, and rinsed with Odyssey Blocking Buffer to remove residual Tween20 prior to scanning for signal using a LI-COR Odyssey imaging system.

**RNA extraction and Northern blotting.** Total RNA was extracted using Trizol (Invitrogen) at 1ml per ~1-2.5 x $10^6$ cells, and subsequently supplemented with 200 µl chloroform. RNA in aqueous phase was precipitated using equal volumes of isopropanol, pelleted, washed in 70% ethanol, air-dried and dissolved in RNase-free water. RNA concentrations were determined using a Nanodrop spectrophotometer (Thermo Fisher Scientific). RNA samples (typically 500 ng) were mixed 1:2 (v/v) with RNA sample loading buffer (Sigma, R4268-5VL), and resolved in a 1.5% denaturing agarose gel containing running buffer (20 mM MOPS, 8 mM sodium acetate, 1 mM EDTA), supplemented with 2% formaldehyde. RNA was transferred to Nylon membranes using overnight passive upward transfer mediated by 10 x SSPE (1.5M NaCl, 0.1M $NaH_2PO_4$, 0.01M EDTA, pH 7.4). Upon transfer, blots were rinsed in 5 x SPPE and crosslinked twice using auto-settings in a UVP CL-1000 UV crosslinker. RNA blots were pre-hybridized in NorthernMAX solution (Ambion) for 1 hour at 42°C, and refreshed with NorthernMAX solution containing 250-500 ng in vitro transcribed biotinylated RNA probe for incubation of a minimum of 3 hours at 42°C. Linearized plasmid containing sequence complementary to FHV RNA1 positions 2719-3007 was used to transcribe biotinylated RNA probes for detection of (+) strand RNAs 1 and 3 on northern blots. Upon hybridization, blots were briefly washed in high-salt buffer (2X SSPE, 0.2% SDS), washed for 20 minutes in low-salt buffer (0.2X SSPE, 0.2% SDS) at 42°C, blocked using Odyssey blocking buffer (LI-COR) containing 1% SDS for 1 hour at room temperature, and incubated in a 1:10,000 dilution of either IRDye680RD-streptavidin or IRDye800CW-streptavidin in blocking buffer containing 1%SDS for 30 minutes at room temperature, washed three times in PBS-0.1% Tween, and once in PBS prior to scanning for signal using a LI-COR Odyssey imaging system.

**Immunofluorescence.** Cells cultured on glass coverslips (Neuvitro GG-18-PDL) were washed three time with PBS (Hyclone, SH30264.01, containing $Ca^{2+}$ and $Mg^{2+}$, for all subsequent washes), fixed in 4% paraformaldehyde in PBS for 10 minutes at room temperature, washed three times in PBS, permeabilized 0.2% Triton X-100 in PBS for 10 minutes, and washed three times for 5 minutes in PBS. Cells were incubated in blocking solution (50 mM $NH_4Cl$, 2% BSA, 0.05% $N_3Na$ in PBS) for 1 hour at room temperature, incubated with primary antibodies diluted in blocking solution for 1 hour at room temperature, washed three times in PBS, incubated for 1 hour at room temperature with secondary antibodies in blocking solution, and washed three times for 5 minutes in PBS. Primary antibodies used were rabbit-anti-FHV protein A (R1194) at 1:500, and mouse-anti-dsRNA (Scicons, clone J2). Secondary antibodies used were goat-anti-rabbit AlexaFluor 568 (ThermoFisher, A11011) and donkey-anti-mouse AlexaFluor 647 (ThermoFisher, A31571), at 1:200 dilution. Coverslips were transferred to glass slides prepared with a drop of NucBlue (Invitrogen, 2 drops diluted in 1 ml PBS) to stain nuclei, incubated for 5 minutes at room temperature, washed three times for 2 minutes in PBS, drained, and mounted using 20 µl ProLong Glass Antifade Mountant (Invitrogen) and cured for at least 1-2 hours in the dark at room temperature. Slides were imaged using either Zeiss ELYRA SIM microscope at the Newcomb Imaging Center at UW-Madison or a wide-field epifluorescence Nikon Ti microscope. Additional image processing used the Fiji (ImageJ) software package.

**Sample preparation for Cryo-electron Tomography.** Mitochondrial isolations were performed using the Qiagen Qproteome mitochondria isolation kit to obtain high-purity mitochondrial fractions. Isolated mitochondria were deposited to 200 mesh Quantifoil cryogrids (2/2) (Cat:

Q2100CR2, Electron Microscopy Sciences) that were glow-discharged for 60s using PELCO easiGlow (Ted Pella, Inc). Grids were plunge-frozen at 22°C using a Vitrobot (FEI, ThermoFisher) under 100% humidity. Fresh blotting paper (Ted Pella, Inc) was applied between different samples. The freezing parameters have been previously described (2). One grid from each frozen group was screened on a Talos 200 keV (ThermoFisher, Inc) equipped with a OneView Gatan camera, the freezing quality was inspected based on the sample density and ice thickness prior to shipping the grids to the Pacific Northwest CryoEM Center (PNCC).

**Cryo-ET imaging acquisition.** Samples were imaged at the PNCC on two Titan Krios microscopes both operated at 300 keV equipped with a Bioquantum K3 direct electron detector. Tilt-series were collected from -54° to 54° at 33K magnification resulting in pixel size 1.653 Å/pixel for the AcR1 sample and 1.645 Å/pixel for the AcRΔ3' sample. The dose rate at K3 camera was 22e$^-$/pixel/sec with a total electron dose of ~180 e-/Å$^2$. A dose symmetric tilt acquisition was used in SerialEM 3.8 and each tilt was fractionated to 20 frames in 0.6 second exposure time at accolated dose rate 4.86 e-/Å2. The target defocus was set at -3.5 µm.

**Cryo-ET Image Processing and Subtomogram averaging.** Image processing for subtomogram averaging was performed as described in (2) and the workflow using *cis*TEM reconstruction (3) and refinement was adapted from (4). Individual tilt-containing frames were motion corrected using Unblur (5). High signal-to-noise ratio tilt-series were manually selected for the following analysis. For *cis*TEM refinement, customized body masks were generated using UCSF Chimera and CryoSparc v3.3.2 (6). For subtomogram averaging of crowns from AcR1-infected cells, ~450 crowns were manually selected from 10 tilt-series to generate an initial model in PEET 1.5 (7). This model served as a template to pick 13104 crowns from 138 tilt-series in emClarity 1.5.3.11 (8). After 3D classification, 3001 crowns were selected for final subtomogram averaging. Initial processing and refinement with no enforced symmetry showed a clear 12-fold symmetry (Fig. S3). Applying C12 symmetry generated the final averaged EM density map with resolution at 17.3 Å. For crowns from AcR1Δ3'-infected cells, an initial model of the particles of interest was generated from a subset of binned tomograms in PEET as described above. Subsequently, 5628 crown-like particles from 34 cryo-tomograms were selected in emClarity and 1685 particles were ultimately used to generate the final average map after 3D classification. The resulting legless crown-like structure, the "proto-crown", showed clear intrinsic 12-fold symmetry. Applying C12 symmetry averaged the final EM density map to a resolution of 12.6 Å (Fig. S3). To compare crowns from FHV-infected cells to the AcR1 and AcR1Δ3' crowns, the same work pipeline was used to re-process our prior published tilt-series (2).

**Sample preparation for single particle analysis.** 5 x $10^8$ Sf9 cells were infected at m.o.i of 20 and shaken at 120 rpm for 60 minutes at room temperature, centrifuged at 500 x g for 5 minutes at room temperature, resuspend in Sf 900 medium at 1.25 x $10^8$ cells in a final volume of 50 ml, and incubated at 28°C for 72 hours. Cells were collected at 500 x g for 5 minutes at room temperature, washed in PBS, and resuspended in 8 ml 0.9% NaCl, centrifuged at 500 x g for 5 minutes at room temperature and resuspended in 2 ml of RSB Hypo Buffer (10 mM HEPES pH 7.5, 10 mM NaCl, 0.2 mM EDTA) containing 1X cOmplete Protease Inhibitor (Sigma, 4693132001). Cells were Incubated on ice for 10 minutes. and dounce-homogenized for 75 passes with a tight pestle. 2.6 ml 2.5X MS Homogenization Buffer (12.5 mM HEPES pH 7.5, 525 mM mannitol, 175 mM sucrose, 2.5 mM EDTA) with 1X cOmplete Protease Inhibitor was added, followed by 3 more dounce passes. Cell lysates were centrifuged at 1,000 x g for 5 minutes at 4°C, and ~6 ml supernatant fraction was saved on ice for later analysis. The lysate pellet fraction was resuspended in 1.3 ml of RSB Hypo Buffer with 1X cOmplete Protease Inhibitor, and subjected to another 75 dounce-passes with a tight pestle. 1.732 ml of 2.5X MS Homogenization Buffer with 1X cOmplete Protease Inhibitor was added followed by 3 more dounce passes and preps were centrifuged at 1,000 x g for 5 minutes at 4°C. Supernatant fractions were centrifuged at 10,000 x g for 10 minutes at 4°C and the crude mitochondria pellet were resuspended to a final concentration of 45 mg/ml in 1X RSB Hypo Buffer supplemented for crosslinking with 2.5 mM disuccinimidyl glutarate (DSG, Thermo Fisher, 20593). Resuspended mitochondria were gently rotated for 30 minutes at 4°C and the crosslinking reaction was quenched using a final concentration of 25 mM Tris pH 8 under gentle rotation for 15 minutes at room temp. An equal volume of 4% n-Dodecyl-B-D-maltoside (DDM) (Gold Biotechnology, DDM5) in Solubilization Buffer (20 mM Tris pH 8.0, 300 mM NaCl, 2 mM $MgCl_2$, 4% glycerol, 2X cOmplete Protease Inhibitor) was added followed by gentle rotation for 30 minutes at room temperature. Preps were centrifuged at 21,000 x g for 15 minutes at 20°C and ~20 ml supernatant fraction was transferred to a new 50 ml conical centrifuge tube. ALFA Selector PE 50% slurry (NanoTag, N-1510) was equilibrated in 4 Bed Volumes (BV) of Wash Buffer (0.05% DDM, 20 mM Tris pH 8.0, 150 mM NaCl, 1% glycerol), and left to settle. Excess wash buffer was removed and equilibrated PE resin was added to the ~20 ml supernatant fraction, gently mixed, and rotated for 1 hour at room temperature.The mixture was loaded onto a Poly-Prep column (Bio-Rad, 7311550, 2 ml bed volume and 10 ml reservoir) and the resin was allowed to pack. The Flow-Through fraction was discarded. The resin was washed with 2.5 ml (12 BV) of Wash Buffer. Wash fractions were discarded, and ALFA-tagged protein A was eluted from PE resin in 428 µl (2 BV) of Elution Buffer (Wash Buffer + 0.02 mM ALFA peptide) by nutating for 25 minutes at room temperature. Samples were collected, and 214 µl (1 BV) of Elution Buffer was added to the resin, nutated again for 25 minutes at room temperature for a second elution. First and second elution were combined (final volume ~ 642 µl). The eluate was concentrated to a final volume of 50-60 µl using a Vivaspin 500 50 kDa MWCO PES column (Cytiva, 28932236) pre-rinsed with 400 µl Washing Buffer. Columns were centrifuged at 2,000 x g for 2 minutes at 4°C, buffer was discarded, eluate was loaded and columns were centrifuged at 2,000 x g for 2 minutes at 4°C. The solution was mixed and centrifuged at 2,400 x g for 2 minutes at 4°C. Solution mixing and centrifugation raised with 400 x g increments was repeated until the volume was between 50-60 µl. Good quality preps usually needed final 3,500-4,000 x g spins. Concentrated eluates were stored at -80°C in Low-Bind microfuge tubes.

**Single particle Cryo-EM sample preparation.** 4µl samples of purified proto-crowns were applied to a 300 mesh Quantifoil cryogrids (1.2/1.3) with a continuous 2nm carbon film (Cat Q3100CR1.3-2nm, Electron Microscopy Sciences, Inc) that were glow-discharged for 7 seconds at 20mA using *GloQube® Plus* glow discharge system (Quorom Inc). The grids were plunge-frozen at 4°C using a Vitrobot Mark IV (FEI, ThermoFisher, Inc) under 100% humidity. Sample was added to the grid and after 30 seconds, the grid was blotted using blot force -15 for 3 seconds. Frozen grids were clipped and then screened on an Artica Talos 200Kv (ThermoFisher, Inc) at PNCC or the UW-Madison CryoEM Center. Freezing quality was assessed based on sample density and ice thickness. High-quality grids were used for data collection on either a Titan Krios equipped with a Bioquantum K3 and a post-column energy filter with a <20eV slit at UW-Madison CryoEM Center on a Titan Krios equipped with a Falcon III or at the PNCC (PNCC proposal ID: EMSLP51742). Detailed imaging parameters are shown in Supplementary Table S1.

**Single Particle Cryo-EM data processing.** For crown-like particles purified from baculovirus-launched expression of Alfa-tagged protein A without a replicable RNA template in Sf9 cells,

9,167 movies were imported into *cis*TEM 2.0 (3) for data processing following the standard *cis*TEM processing workflow (Fig. S4). After motion correction and contrast transfer function (CTF) parameter estimation, only micrographs with a detected CTF fit resolution better than 3.5 Å were kept for further processing. Initially ~790k particles were automatically picked using the *cis*TEM disc picker and subjected to 2D classification. The resulting 2D class averages clearly revealed a mixture of 11- and 12-fold symmetric particles, which exhibited a strong preference for the top view orientation with around ~22k particles classifying into side views, ~20k particles classifying into intermediate views and ~330k particles classifying into top views. For efficiency, the initial round of processing was carried out using only side views. An ab initio from the best selected class averages performed without symmetry resulted in a reconstruction which demonstrated clear C11 symmetry. Auto-refinement with C11 symmetry imposed resulted in an initial 3.9 Å resolution structure. 3D classification with C11 resulted in the 3D classes which exhibited heterogeneity in the polymerase regions, and also identified two subsets one "larger diameter" and one "smaller diameter" which we reasoned to be separation of the C11 and C12 symmetries. The best smaller diameter classes were combined (~14k particles) and further refined with C11 to obtain a map with a ~ 3.1 Å global resolution. This process was then repeated with C12 symmetry imposed with the best larger diameter classes being combined (~9k particles), and being further refined with C12 symmetry resulting in a ~3.2 Å global resolution structure. The final reconstructions were created by adding approximately 10% top views which were picked based on the 2D classification as either C11 or C12, resulting in maps with the same global resolution but slightly improved density as viewed from the top. Inclusion of more top views increased the global resolution, but led to reconstructions which were stretched from the side and more difficult to interpret. In both cases, the highest resolution used for refinement was 4.5 Å. The workflow and data acquisition parameters are shown in Table S1.

For crown-like particles (CLP) from FHV-alfa-infected S2 cells, 11,629 movies were imported into *cis*TEM, motion corrected and CTF parameter estimated. Following automatic picking, ~1.3 million particles were selected and subjected to 2D classification. The 2D class averages once again demonstrated a severe preference for top views, however all of the observed class averages contained 12 subunits (Fig. S5), suggesting that in this dataset the C12 form was dominant. The data acquisition parameters are shown in Table S1.

**Model building of FHV protein A.** We built the FHV protein A atomic model starting from the N-terminus due to its related high-resolution EM densities that appear in both C11 and C12-maps. *De novo* building of alfa-tagged FHV protein A was initiated by fitting the highest ranked Alphafold2 (9) predicted model from COLABFOLD (10) into both C11- and C12-sharpened maps in Coot (11). Higher resolution C11-map allowed us to affirmatively assign residues of FHV protein A 55-396 with gaps between residues 123-150. We did not observe clear density for the extreme N-terminal residues from 1 to 54. The built model was then iteratively refined with Phenix v1.20 (12) and Isolde v1.3 (13) in ChimeraX v1.3 (14). The validation of the model used MolProbity (15), and validation tools in Phenix. Ligand-binding locations for SAH and GTP were calculated and simulated using Autodock Vina v1.20 in UCSF Chimera (16). Refinement statistics are shown in Table S2.

**Polymerase Sharpening.** To improve the resolution of the FHV Pol domain we carried out symmetry expansion with density subtraction in *cis*TEM to isolate pairs of polymerase domains (3, 17). A mask was manually created in UCSF Chimera which contained two adjacent asymmetric subunits from the C11 imposed structure. Using the *cis*TEM command *symmetry_expand_stack_and_par,* this mask was used, in combination with C11 symmetry to create a subtracted dataset containing isolated and centered polymerase pairs (3, 17). These pairs were then further refined by local refinement and 3D classification in *cis*TEM, with the best 3D class kept as the final refined Pol density.

To generate a partial backbone trace of Pol we first used UCSF Chimera to fit the structure predicted by AlphaFold2 (9) into the refined Pol EM density. Visual inspection showed good agreement for most of the structure with only minor offsets between EM density and predicted model. Regions that were unclear in the EM density or differed between the EM density and model were removed (aa 487-533, 630-53, 816-28 and C-terminal aa 862-96) and the remaining model, covering 80% of the Pol domain to residue 861, was refined using Phenix to correct residual minor offsets to produce the partial backbone trace model of the Pol domain.

### Supplementary Figures

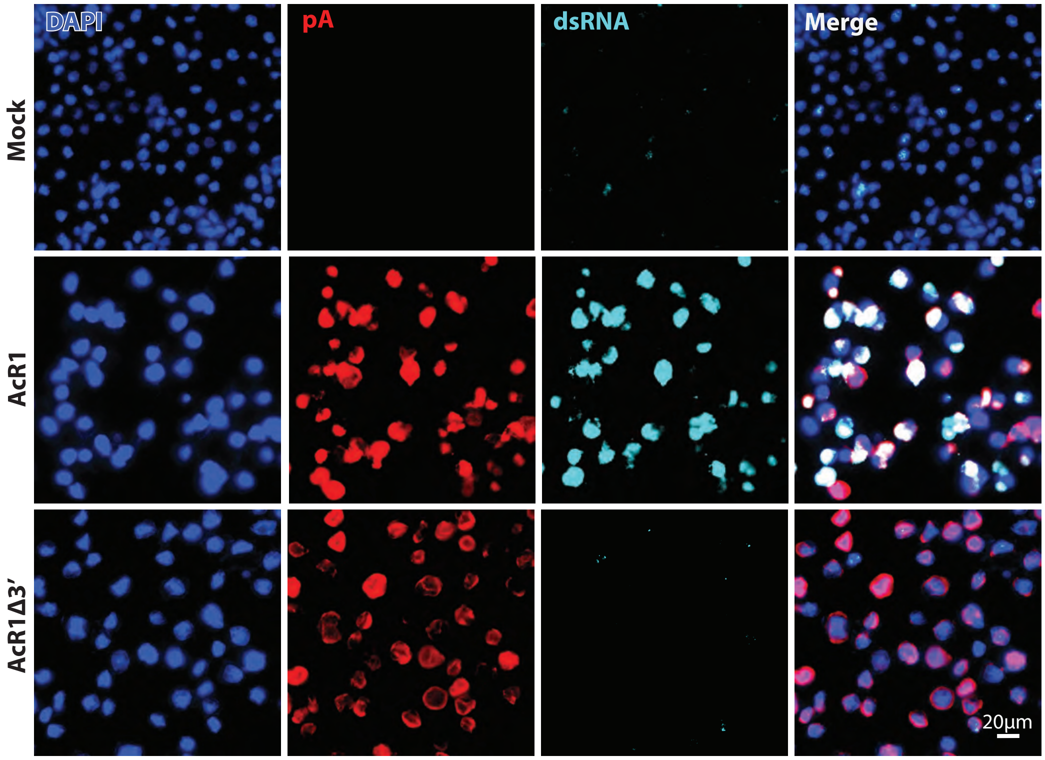


**Figure S1.** Immunofluorescence analysis of baculovirus-launched nodavirus RNA1 replication. Nodavirus protein A was expressed in S. frugiperda Sf9 cells infected with baculoviruses AcR1 or AcR1Δ3′ at an m.o.i. of 20 at 48 hours post-infection. However, dsRNA was only detected in AcR1-infected cells, consistent with RNA1 replication dependent on protein A recognition of 3′ UTR sequences present in AcR1-expressed wt RNA1 but not in the 3′UTR-deleted RNA1 derivative expressed by AcR1Δ3′.

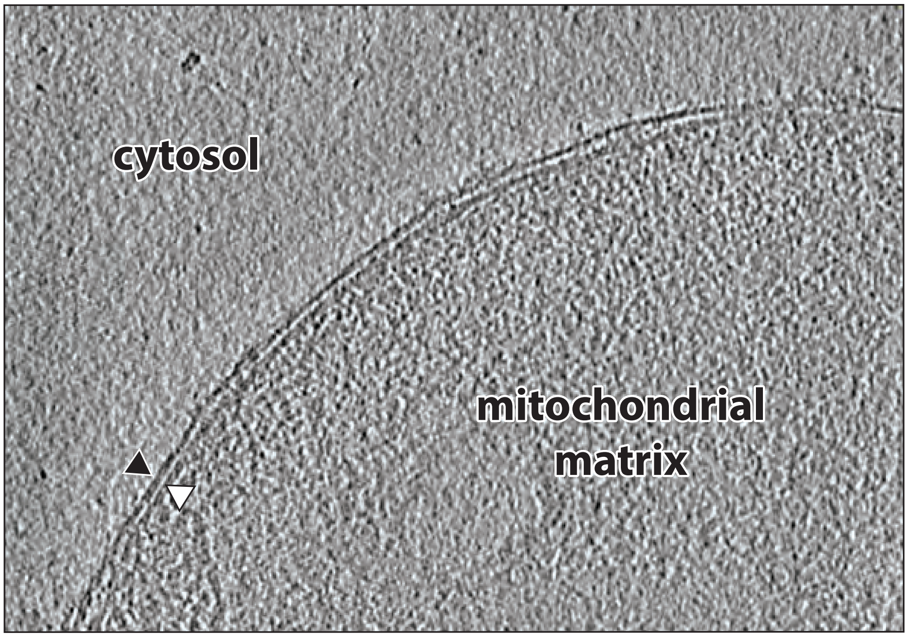


**Figure S2.** Cryo-ET imaging shows that healthy mitochondria from uninfected Sf9 cells lack the spherular invaginations of the outer mitochondrial membrane (OMM) and the cytosol-facing crown-like densities associated with nodavirus RNA replication complexes. The tightly opposed outer and inner mitochondrial membranes are indicated by a black and a white arrowhead, respectively.

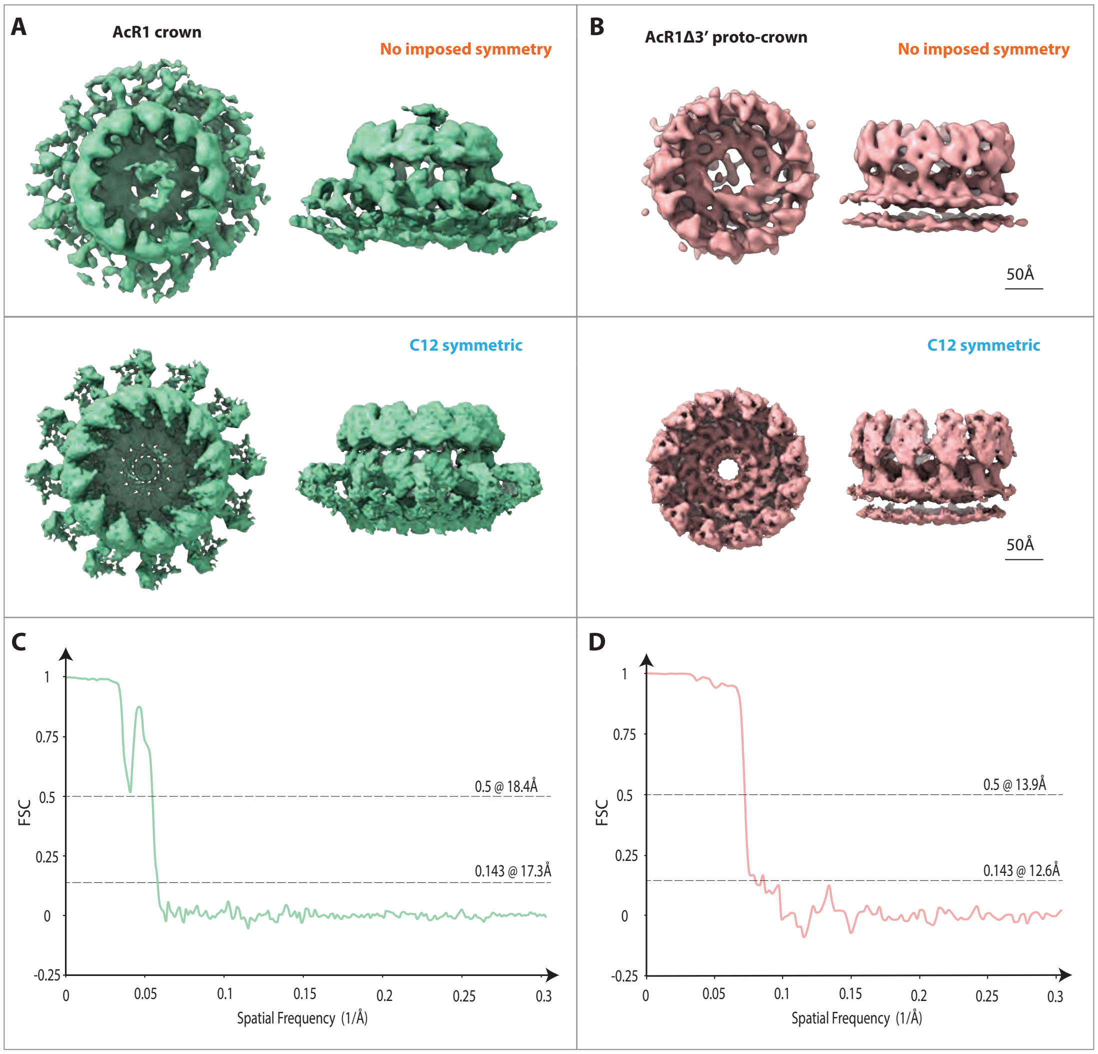


**Figure S3.** Top and side views of **(A)** mature, RNA-replication active crowns (green) from baculovirus AcR1-infected Sf9 cells and **(B)** proto-crowns (pink) from AcR1Δ3′-infected Sf9 cells, as imaged by subtomogram- averaged cryo-ET with no imposed symmetry (upper panels) and with C12 symmetry imposed (lower panels).
(C - D) Fourier Shell Correlation curves with FSC 0.5 and 0.143 threshold values indicating the resolution of the C12 symmetry-imposed structures for AcR1-induced crowns and AcR1Δ3′-induced proto-crowns, respectively.

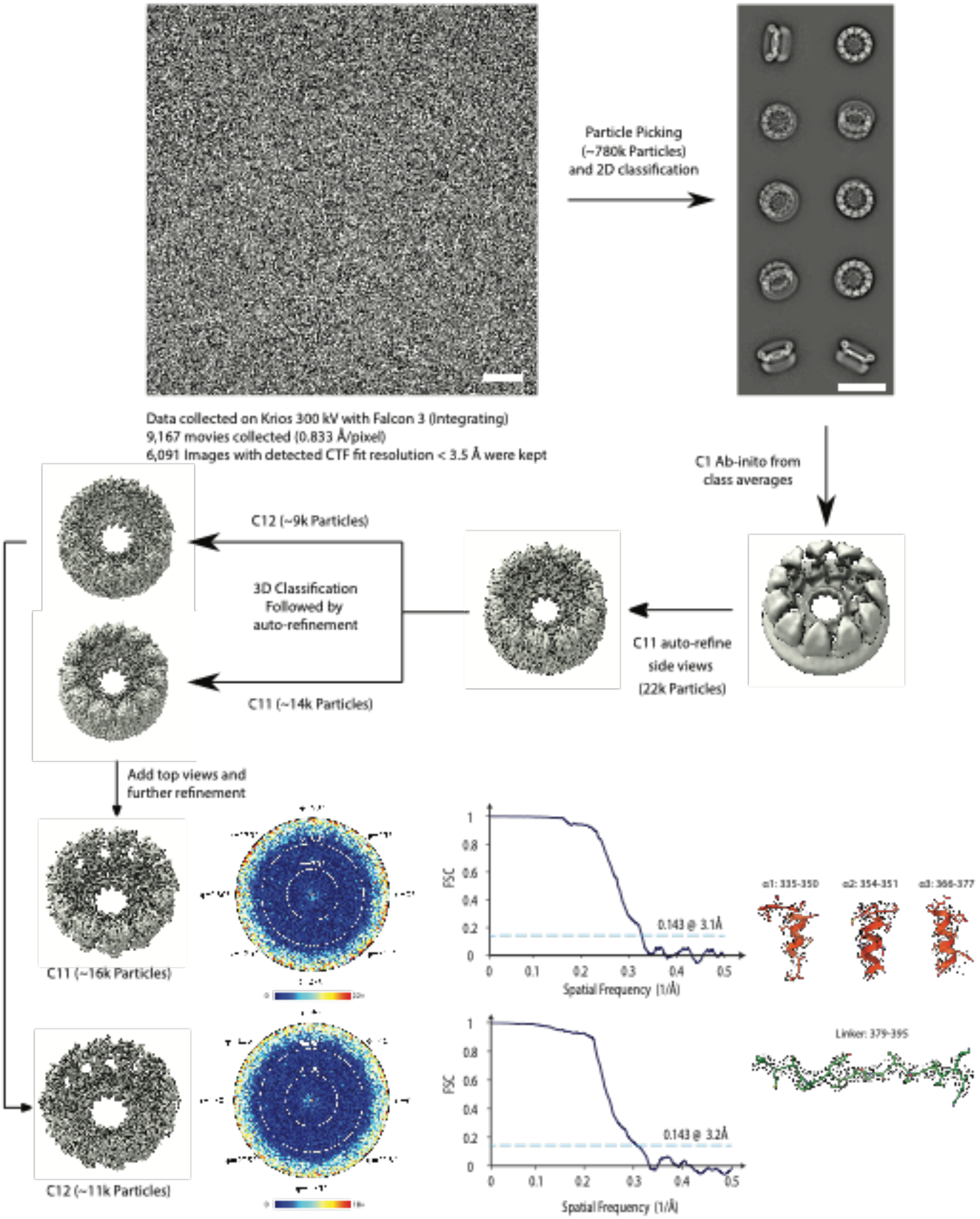


**Figure S4.** Graphical description of single particle cryo-EM processing methods used in this study, as described in the Supplementary Methods. The workflow illustrates a typical cryo-EM image and representative class averages (scale bars = 35 nm). Final reconstructions of AcR1Δ3'-induced baculovirus C11 and C12 proto-crowns from 3D classification are shown alongside the angular plot demonstrating the distribution of particle views, the Fourier shell correlation used for the global resolution estimation and examples of protein structural model building within the refined cryo-EM density.

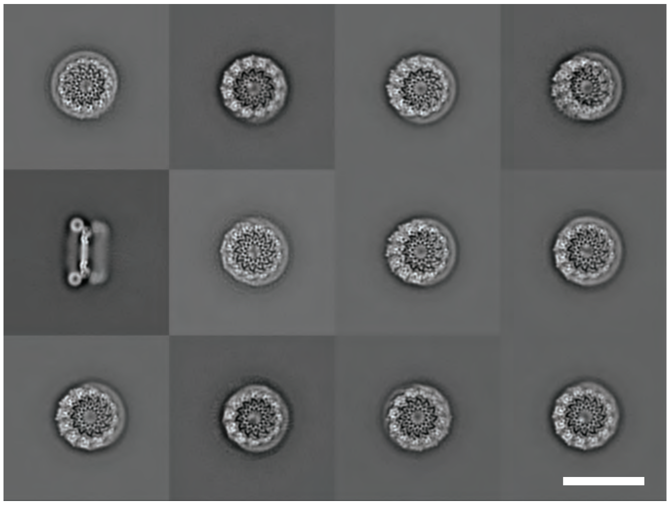

**Figure S5.** 2D classes of proto-crown particles purified from S2 cells infected with virions of FHV-alfa, a fully infectious FHV derivative engineered to add an alfa tag to protein A's C-terminus.  Scale bar = 35 nm.

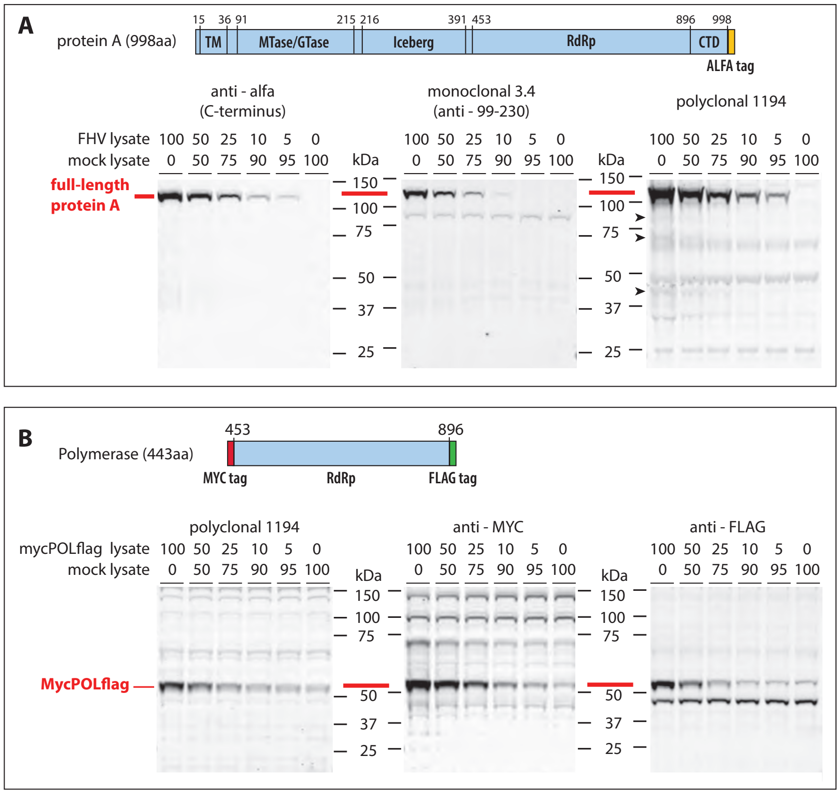


**Figure S6.** Nodavirus protein A does not undergo post-translational processing to release an RNA polymerase unit-sized proteolytic product.

**(A)** Western blots of protein A in S2 cells infected with FHV-alfa, a fully infectious FHV derivative engineered to fuse an alfa tag to protein A's C-terminus. A monoclonal antibody against N-proximal protein A aa 99-230, a polyclonal antiserum against protein A that recognizes the protein A polymerase domain (see panel B), and a fluorescent nanobody against the C-terminal alfa epitope tag all detect essentially only full-length protein A in lysates from FHV-infected cells. Identification of host protein background in the westerns is facilitated by progressive dilution of the infected lysate in a mock-infected control lysate to dilute virus-specific proteins but not host proteins. The polyclonal antiserum does reveal a few smaller protein A-specific proteins (arrowheads), but their minimal amounts are vastly insufficient to account for the 12 polymerase units in the upper rings of mature crown-structures, which comprise half of all FHV polymerase in infected cells.

**(B)** Western blots confirming that polyclonal antiserum 1194 used in panel (A) recognizes epitopes in the FHV protein A RNA polymerase domain expressed without other viral sequences, ruling out the possibility that the serum would have failed to detect any such "free" polymerase in panel (A).

### Supplementary Tables

**Supplementary Table S1** Cryo-EM data acquisition parameters

| Data collection and Processing | FHV A-alfa infection | Baculovirus-launched FHV protein A-alfa |
|---|---|---|
| Magnification | 81 K | 96 K |
| Voltage (keV) | 300 | 300 |
| Total Electron Dose (-e/Å2) | 50 | 100 |
| Dose fractionations (frames) | 55 | 63 |
| Defocus range (µm) | 0.5-2.0 | 0.5-2.0 |
| Detector | Gatan K3 Summit | Falcon III Integrating |
| Energy filter | Gatan GIF Quantum 20eV slit | NA |
| Raw Pixel Size (Å) | 1.064 | 0.833 |
| Super Resolution Pixel Size (Å) | 0.532 | NA |
| Number of collected movies | 11,629 | 9,167 |
| Number of selected micrographs | N/A | 6,091 |
| Software | SerialEM 3.8 | SerialEM 3.8 |

**Supplementary Table S2** Refinement and validation statistics

| | FHV Protein A Floor C12 ring (EMDB-XXXXX) | Protein A Floor C11 ring (EMDB-XXXXX) |
|---|---|---|
| **3D Reconstruction** | | |
| Software | *cis* TEM | *cis* TEM |
| Number of selected particles | 11,093 | 13,701 |
| Map resolution (Å) | 3.2 | 3.1 |
| Symmetry imposed | C12 | C11 |
| FSC threshold | 0.143 | 0.143 |
| **Refinement** | | |
| Initial model used | Alphafold prediction | Alphafold prediction |
| Model build | *De novo* | *De novo* |
| **Model compostion** | | |
| Non-hydrogen atoms | 29916 | 28820 |
| Protein residues | 3696 | 3564 |
| Chains | A-L | A-K |
| **Bonds (RMSD)** | | |
| Bonds length (Å) (# > 4σ ) | 0.002 | 0.003 |
| Bonds Angle (º) (# > 4σ ) | 0.452 | 0.683 |
| min/max/mean | 110.01/205.59/143.34 | 50/146.44/92.41 |
| **Validation** | | |
| Molprobity score | 1.68 | 1.58 |
| Clash score | 15.12 | 13.51 |
| **Ramachandran plot statistics (%)** | | |
| Favored | 98.36 | 97.5 |
| Allowed | 1.64 | 2.5 |
| Outlier | 0 | 0 |

**SI References**


1. P. A. Venter, N. K. Krishna, A. Schneemann, Capsid protein synthesis from replicating RNA directs specific packaging of the genome of a multipartite, positive-strand RNA virus. *J. Virol.* **79**, 6239–6248 (2005).
2. N. Unchwaniwala, *et al.*, Subdomain cryo-EM structure of nodaviral replication protein A crown complex provides mechanistic insights into RNA genome replication. *Proc. Natl. Acad. Sci. U. S. A.* **117**, 18680–18691 (2020).
3. T. Grant, A. Rohou, N. Grigorieff, cisTEM, user-friendly software for single-particle image processing. *eLife* **7** (2018).
4. T. Ni, *et al.*, High-resolution in situ structure determination by cryo-electron tomography and subtomogram averaging using emClarity. *Nat. Protoc.* **17**, 421–444 (2022).
5. T. Grant, N. Grigorieff, Measuring the optimal exposure for single particle cryo-EM using a 2.6 Å reconstruction of rotavirus VP6. *eLife* **4**, e06980 (2015).
6. A. Punjani, J. L. Rubinstein, D. J. Fleet, M. A. Brubaker, cryoSPARC: algorithms for rapid unsupervised cryo-EM structure determination. *Nat. Methods* **14**, 290–296 (2017).
7. J. M. Heumann, A. Hoenger, D. N. Mastronarde, Clustering and variance maps for cryo-electron tomography using wedge-masked differences. *J. Struct. Biol.* **175**, 288–299 (2011).
8. B. A. Himes, P. Zhang, emClarity: software for high-resolution cryo-electron tomography and subtomogram averaging. *Nat. Methods* **15**, 955–961 (2018).
9. J. Jumper, *et al.*, Highly accurate protein structure prediction with AlphaFold. *Nature* **596**, 583–589 (2021).
10. M. Mirdita, *et al.*, ColabFold: making protein folding accessible to all. *Nat. Methods* **19**, 679–682 (2022).
11. P. Emsley, B. Lohkamp, W. G. Scott, K. Cowtan, Features and development of Coot. *Acta Crystallogr. D Biol. Crystallogr.* **66**, 486–501 (2010).
12. D. Liebschner, *et al.*, Macromolecular structure determination using X-rays, neutrons and electrons: recent developments in Phenix. *Acta Crystallogr. Sect. Struct. Biol.* **75**, 861–877 (2019).
13. T. I. Croll, ISOLDE: a physically realistic environment for model building into low-resolution electron-density maps. *Acta Crystallogr. Sect. Struct. Biol.* **74**, 519–530 (2018).
14. E. F. Pettersen, *et al.*, UCSF ChimeraX: Structure visualization for researchers, educators, and developers. *Protein Sci. Publ. Protein Soc.* **30**, 70–82 (2021).
15. C. J. Williams, *et al.*, MolProbity: More and better reference data for improved all-atom structure validation. *Protein Sci. Publ. Protein Soc.* **27**, 293–315 (2018).
16. J. Eberhardt, D. Santos-Martins, A. F. Tillack, S. Forli, AutoDock Vina 1.2.0: New Docking Methods, Expanded Force Field, and Python Bindings. *J. Chem. Inf. Model.* **61**, 3891–3898 (2021).
17. J. Jung, *et al.*, High-resolution cryo-EM structures of outbreak strain human norovirus shells reveal size variations. *Proc. Natl. Acad. Sci. U. S. A.* **116**, 12828–12832 (2019).
